# MRGM: An enhanced catalog of mouse gut microbial genomes substantially broadening taxonomic and functional landscapes

**DOI:** 10.1101/2024.08.12.607507

**Authors:** Nayeon Kim, Chan Yeong Kim, Junyeong Ma, Sunmo Yang, Dongjin Park, Sang-Jun Ha, Peter Belenky, Insuk Lee

## Abstract

Mouse gut microbiome research is pivotal for understanding the human gut microbiome, providing insights into disease modeling, host-microbe interactions, and the dietary influence on the gut microbiome. To enhance the translational value of mouse gut microbiome studies, we need detailed and high-quality catalogs of mouse gut microbial genomes. We introduce the Mouse Reference Gut Microbiome (MRGM), a comprehensive catalog with 42,245 non-redundant mouse gut bacterial genomes across 1,524 species. MRGM marks a 40% increase in the known taxonomic diversity of mouse gut microbes, capturing previously underrepresented lineages through refined genome quality assessment techniques. MRGM not only broadens the taxonomic landscape but also enriches the functional landscape of the mouse gut microbiome. Using deep learning, we have elevated the Gene Ontology annotation rate for mouse gut microbial proteins from 3.2% with orthology to 60%, marking an over 18-fold increase. MRGM supports both DNA- and marker-based taxonomic profiling by providing custom databases, surpassing previous catalogs in performance. Finally, taxonomic and functional comparisons between human and mouse gut microbiota reveal diet-driven divergences in their taxonomic composition and functional enrichment. Overall, our study highlights the value of high-quality microbial genome catalogs in advancing our understanding of the co-evolution between gut microbes and their host.

## Introduction

The past decades have shown a strong link between the gut microbiome and human diseases, leading to the concept of microbiome medicine ^1^. This approach considers the gut microbiome as a potential biomarker and therapeutic target ^2–4^. Research often begins by comparing the microbiomes of patients and healthy individuals, using reference microbial genomes to identify disease-associated microbial taxa. The quality of these reference genomes is crucial for successful discovery in such studies. Consequently, there is an increasing effort to develop comprehensive and high-quality reference microbial genomes through metagenome-assembled genome (MAG) collection to enhance our understanding of the taxonomic and functional landscape of the gut microbiome, which remains mostly unexplored ^5–10^.

Mouse models have been pivotal in researching human diseases, including playing crucial roles in the study of the human gut microbiome. These models allow for the examination of complex interactions between gut microbiota and the host immune system, metabolism, and genetics under controlled conditions. The impact of gut microbiota on various diseases, such as inflammatory bowel disease (IBD), obesity, diabetes, autoimmune diseases, and cancer, has been explored using mouse models. Furthermore, the effects of drugs or diets on the gut microbiome have also been studied extensively in these models. However, translating insights from mouse studies to human applications faces challenges due to significant differences in gut microbiota ^11, 12^. Consequently, a comprehensive understanding of the taxonomic and functional landscape of the mouse gut microbiome is essential for advancing human gut microbiome research.

The cataloging of mouse intestinal microbial genomes through MAGs has seen significant advancements, with several catalogs now published ^13–16^. However, these catalogs predominantly feature medium quality (MQ) MAGs, defined by a completeness of ≥50% and contamination of ≤5%. Such MQ MAGs are missing a considerable portion of core genes, leading to a substantial loss of core gene information. This loss can severely distort the downstream functional analysis of species’ pangenomes ^17^. Moreover, microbial species clusters composed solely of MQ genomes may, in fact, be nonexistent, raising concerns about the accuracy of these catalogs. Therefore, it is recommended to prioritize the analysis of microbiomes using catalogs that contain only complete genomes and near-complete (NC) MAGs, defined by a completeness of ≥90% and contamination of ≤5%. The recently released catalogs— integrated mouse gut metagenome catalog (iMGMC) ^14^, mouse gut bacterial catalog (MGBC) ^15^, and comprehensive mouse microbiota genome (CMMG) catalog ^16^— comprise 9,053, 26,640, and 22,765 NC genomes, corresponding to 830, 1,094, and 1,179 bacterial species, respectively. This achievement marks a significant step forward in understanding the mouse gut microbiome. Despite these advancements, a substantial portion of the mouse gut microbiome remains unexplored, suggesting that further research is needed to achieve a more comprehensive cataloging of its microbial genomes.

Herein, we introduce the mouse reference gut microbiome (MRGM), featuring 42,245 non-redundant NC genomes that encompass 1,524 bacterial species, including 473 novel mouse gut microbial species with high-quality genomes. This addition expands the high-quality taxonomic landscape of the mouse gut microbiome by more than 40%. Additionally, MRGM facilitates taxonomic profiling using both DNA- and marker-based methods supported by custom databases. Through analyses of both simulated and real metagenome sequencing data, the superiority of MRGM in the accuracy of taxonomic profiling with these methods has been established. Moreover, we significantly broadened the functional landscape of the mouse gut microbiota by employing a deep learning model, which boosted Gene Ontology (GO) annotation rate from approximately 3.2% via orthology to around 60% of gut microbial protein families in mice, achieving a more than 18-fold increase. Comparative analyses of gut microbial genome catalogs between humans and mice highlighted the taxonomic and functional divergences in their gut microbiomes, underscoring differences related to their distinct dietary patterns.

## Materials and Methods

### Whole metagenome sequencing of mouse fecal samples

We conducted WMS on fecal samples from 40 C57BL/6J mice. Initially, we subjected 100 ng of DNA from each sample to fragmentation using a Covaris LE200 Focused-ultrasonicator, creating blunt-ended DNA fragments. We then selected DNA fragments of approximately 350 bp in length. These fragments underwent ligation with indexing adapters and were subsequently amplified through eight cycles of PCR. The quality and size distribution of the resulting libraries were assessed using D1000 Screen Tapes on a Tapestation 4200, both from Agilent Technologies. Library preparation followed the TruSeq Nano DNA Library Preparation Guide’s protocols. We quantified the libraries using KAPA Library Quantification kits, specifically designed for Illumina sequencing platforms, as detailed in the qPCR Quantification Protocol Guide by KAPA BIOSYSTEMS (catalog #KK4854). Sequencing was carried out on the HiSeq 4000 system by Illumina, based in San Diego, USA, targeting a sequencing depth between 12 and 47 Gbp.

### Metagenome *de novo* assembly and binning

In our study, we supplemented in-house WMS data from 40 mouse fecal samples with publicly available WMS data for 1,837 samples that were recently made available but not yet incorporated into existing catalogs. We excluded samples from mice that had undergone fecal microbiota transplantation from humans. Sequence reads were cleaned of adapters and low-quality bases using Trimmomatic v0.39 ^18^, employing settings ‘ILLUMINACLIP:adapter sequence:2:30:7 LEADING:5 TRAILING:5 SLIDINGWINDOW:4:5 MINLEN:36’.

Subsequently, we eliminated sequences contaminated by the host genome using Bowtie2 v2.2.5 ^19^, aligning reads against the mouse genome GRCm39 (GenBank assembly accession GCA_000001635.9). For assembling high-quality reads into contigs, we primarily used MetaSPAdes v3.15.3 ^20^, with MEGAHIT v1.2.9 ^21^ serving as an alternative for instances where MetaSPAdes failed or when dealing with single-end sequencing data. Binning, the process of organizing contigs into genome bins, was carried out through an ensemble approach. This process began with aligning sequence reads back to the contigs using Bowtie2. We then employed three separate binning tools— MetaBAT2 v2.12.1 ^22^, MaxBIN2 v2.2.7 ^23^, and CONCOCT v1.1.0 ^24^—to create genome bins independently, setting the minimum contig length at 1,000 for MaxBin2 and CONCOCT, and 1,500 for MetaBAT2. The binning results were integrated using the bin refinement module of MetaWRAP v1.3.2 ^25^. To evaluate the completeness and contamination of genome bins, we utilized CheckM2 v0.1.2 ^26^. Additionally, to compare the completeness estimates provided by CheckM and CheckM2 for novel lineages, we employed the lineage-specific workflow available in CheckM v1.1.3 ^27^. Notably, we adapted MetaWRAP code to select the genome bin of the highest quality based on CheckM2 v0.1.2 evaluations, replacing the older CheckM assessment. Further assessment was conducted with GUNC v1.0.5, a tool designed to detect genome chimerism and calculate clade separation scores (CSS) ^28^. Genomes displaying a CSS greater than 0.45 were excluded, adhering to the default threshold set by GUNC.

### Public MAGs and isolated genomes in the MRGM catalog

For the MRGM catalog, we sourced publicly available MAGs and isolated genomes pertinent to the mouse gut microbiome from the MGBC ^15^. Additionally, we included isolated genomes from the expanded mouse intestinal bacterial collection (miBC) ^29^. Single-cell amplified genomes (SAGs) were obtained from assembled genome sequences as documented in previous studies ^30^.

### Assessment of sequencing depth effect on MAG assembly

To evaluate whether increased sequencing depth improves MAG assembly, we generated simulated datasets across various sequencing depths. We selected 10 WMS samples with the largest sequencing depths from a pool of 40 samples, each sequenced to ultra-deep levels (≥40 Gbp or ≥130 million read pairs). For each sample, we randomly selected 0.5, 2.5, 5, 10, 20, 40, 80, and 125 million read pairs, corresponding to total sequencing depths of 150 Mbp, 750 Mbp, 1.5 Gbp, 3 Gbp, 6 Gbp, 12 Gbp, and 37.5 Gbp for a read pair length of 300 bp. In total, 80 simulated datasets (10 samples × 8 depths) were assembled using the same in-house pipeline. To determine whether increased sequencing depth improved the quality of conspecific genomes, we performed average-linkage hierarchical clustering at a Mash distance cutoff of 0.1 (equivalent to Mash ANI 90%). If MAGs from the same sample but different sequencing depths clustered together, they were considered conspecific genomes. We then compared the quality of these conspecific MAGs across adjacent sequencing depths to assess improvements.

### Generation of species clusters

To delineate species clusters, we adopted a two-tiered iterative clustering approach, consisting of an initial fast clustering phase followed by a more precise secondary clustering. The preliminary clustering utilized average-linkage hierarchical clustering with a threshold of 0.2, employing Mash v2.3 ^31^ to compute the distance based on the taxonomic rank of order as annotated by GTDB-tk v2.1.1 ^32^. For this purpose, Mash calculated all pairwise genome distances using a sketch size of 10,000, enabling rapid distance estimation. However, it is important to note that Mash’s accuracy may decline with lower genome coverage. To enhance the precision of the initial clusters, we performed refined secondary clustering utilizing ANImf metrics from dRep ^33^. ANImf scores were computed for every genome pair within each preliminary cluster. To mitigate potential overestimations of average nucleotide identity (ANI) resulting from local alignments, we applied a minimum coverage threshold of 0.6 for each comparison ^34, 35^. Clustering at this stage was based on an ANI threshold of ≥95%, indicating that genomes within the same cluster are considered to belong to the same species. Within each cluster, we selected a representative genome using a genome intactness score formula *S* = *Completeness* − 5 × *Contamination* + 0.5 × *N*50 , prioritizing the genome with the highest score. For the enumeration of conspecific genomes within each species cluster, we employed average-linkage hierarchical clustering with a cutoff of 99.9% ANI and 0.81 coverage, based on the ANI values obtained during the species clustering process. The increase in coverage cutoff to 0.81 was justified by our focus on NC genomes, which assured a minimum completeness of 90%. This adjustment to the coverage cutoff, derived from the product of the minimum completeness values of two paired genomes.

### Taxonomic annotation of species using GTDB

For the taxonomic annotation of species, we utilized representative genomes and classified them according to the GTDB r207 ^36^. Using GTDB-Tk v2.1.1 ^32^, we assigned taxonomic classifications to the query genomes based on GTDB reference phylogenetic tree, which utilizes 120 bacterial and 122 archaeal marker genes. GTDB-Tk conducted an alignment of these marker genes and produced multiple sequence alignments for each representative genome of the species. We then constructed a phylogenetic tree using IQ-TREE v2.2.0.3 ^37^, based on the multiple sequence alignments of the concatenated sequences of the 120 bacterial marker genes. For the visualization of the resulting phylogenetic tree, we employed the Interactive Tree of Life (iTOL) v6 ^38^.

### 16S rRNA sequence prediction from bacterial species genomes

We utilized barrnap v0.9 ^39^ to predict 16S rRNA sequences from the genomes of bacterial species, adjusting the e-value threshold from the default 1e-06 to 1e-04 to increase sensitivity. Given the high conservation of rRNA genes and the challenges in predicting them from MAGs derived from short-read sequences, we expanded our analysis to include all redundant genomes, not just the representative ones. For species whose 16S rRNA sequences could not be predicted, we obtained genome sequences for the corresponding species from the GTDB r207 ^36^, enabling the prediction of 16S rRNA sequences for an additional 69 species.

### Surveying genera prevalence using kraken2 and bracken

To assess the prevalence of genera, we estimated the relative abundance of 1,524 species in the MRGM catalog across 1,877 WMS samples (1,837 public and 40 in-house samples) using kraken2 v2.1.2 ^40^, coupled with bracken v2.8 ^41^. To minimize false positive identifications, we configured kraken2 with the settings ‘--confidence 0.2 --minimum-hit-groups 3’. A genus was deemed present in a sample only if its relative abundance surpassed the threshold of 1e-6. We excluded 34 samples with shallow sequencing (<100Mbp) from the survey report, resulting in genera prevalence being reported across 1,843 samples.

### Cataloging mouse gut microbial proteins and functional annotation

To compile a comprehensive catalog of mouse gut microbial proteins, we predicted coding sequences (CDS) from genomes using Prokka v1.14.6 ^42^ with the parameter “--kingdom Bacteria". To minimize redundancy in the protein sequences, we employed linclust from MMseq2 v13.45111 ^43^, using the parameters "--cov-mode 1 --c 0.8 --cluster-mode 2 --kmer-per-seq 80 --min-seq-id 1" for clustering at 100% sequence similarity. This process was then sequentially repeated for clustering at 90%, 80%, and 50% sequence similarity, adjusting the --min-seq-id parameter to 0.90, 0.80, and 0.50, respectively. The protein families with 90% similarity were annotated using eggNOG-mapper v2.1.2 ^44, 45^ with the parameter "-m diamond" for functional annotations, including protein families (PFAMs) ^46^, KEGG pathways/modules/orthologs ^47^, Gene Ontology (GO) ^48^, Carbohydrate-Active enZymes (CAZy) terms ^49^, and enzyme commission (EC) numbers ^50^.

Given the typically low annotation rate of GO terms by eggNOG-mapper, we supplemented our approach with deepGOPlus v1.0.1 ^51^ with threshold 0.3 to predict GO functions more comprehensively. We focused exclusively on bacteria-associated GO terms to avoid false-positive annotations. From the UniRef90 database ^52^, updated in September 2023, we filtered to identify only those protein families derived from bacterial organisms. Bacteria-associated GO terms were then determined based on the GO annotations present in these filtered UniRef90 bacterial protein families. We noted that the GO cellular component ’cellular anatomical entity (GO:0110165)’ was annotated for over 90% of protein families by deepGOPlus. Given its broad definition and absence in eggNOG-mapper assignments, this likely indicates a technical bias in deepGOPlus. Therefore, we excluded this GO term from our protein functional annotation.

To facilitate a comparative analysis between the annotations provided by eggNOG-mapper and deepGOPlus, we applied identical filtering criteria for GO terms predictions with both tools, ensuring that only bacteria-relevant terms were considered. Subsequently, we assessed the functional diversity across the 1,524 species represented in the MRGM catalog by calculating the Jaccard distance based on their GO term annotations.

### Comparison of functional annotations: deepGOPlus vs. eggNOG-mapper

To compare functional annotations, we began by predicting GO terms for protein families with 90% similarity using either eggNOG-mapper v2.1.2 ^44, 45^ or deepGOPlus v1.0.1 ^51^. We then profiled 1,524 representative genomes from each species based on GO terms derived from each method. Following this, we assessed the functional distance between genomes utilizing the Jaccard distance metric and performed agglomerative clustering on the genomes based on these functional distances. We evaluated the homogeneity of taxonomic classes within clusters by measuring the consistency of genus labels among the genomes in each cluster, for example, when assessing at the genus level. This evaluation was performed at different distance thresholds, with the number of clusters varied from 1 to 200. Higher homogeneity within genome clusters suggests superior performance of the functional annotation method for the encoded proteins.

### Construction of pan-genomes and their size comparison across MRGM species

For 1,120 of the 1,524 species cataloged in the MRGM, each featuring more than one non-redundant genome, we constructed pan-genomes using panaroo v1.2.9 with the parameter ‘--clean-mode strict’ ^53^. The input for panaroo comprised GFF files generated by Prokka, facilitating the pan-genome analysis. The gene presence-absence matrix produced by panaroo, detailing the occurrence of genes across genomes, served as the basis for this comparison. We conducted a rarefaction and extrapolation analysis on this matrix using the iNEXT function, with the data type set to ‘incidence raw’ ^54^, to evaluate and compare the expansiveness of the pan-genomes across these selected species.

### Construction of a custom database of species-specific marker genes for MRGM

We employed methodologies akin to those used in MetaPhlAn4 v4.0.2 ^55^ to extract unique marker genes across the 1,524 species identified within the MRGM catalog. Initially, we identified potential marker genes using panaroo-generated pangenomes for 1,120 species with multiple conspecific genomes. For species represented by a single genome, all protein sequences within that genome were considered for pangenome construction. We then clustered these protein sequences into families at 90% similarity using Linclust, allowing us to assess the ’coreness’ (the proportion of conspecific genomes containing a given protein family) and ’uniqueness’ (the exclusivity of a gene family to a limited number of species) for each protein family. Only protein sequences ranging from 150 to 1,500 amino acids and having less than 1.5 copies per genome on average were considered to avoid paralogous sequences. We regarded gene families satisfying specific thresholds for coreness and uniqueness as marker gene candidates, with a maximum of 800 gene families per species selected. We applied a coreness threshold of 50% for species with over 100 conspecific genomes and 60% for those with fewer conspecific genomes, following the same thresholding parameters used in the MetaPhlAn4 pipeline. The uniqueness criterion was established at a maximum of 10, meaning that gene families found in no more than 10 species were considered unique enough to be retained.

In the subsequent step, we refined our selection of marker genes from these candidates. We fragmented the DNA sequences of these genes into 150 bp segments and aligned them against the 42,245 conspecific genomes using Bowtie2 with the ’--very-sensitive’ option. This alignment process helped reassess the coreness and uniqueness values for each marker gene candidate. The coreness threshold was set at 60%, and the uniqueness threshold at 1, indicating the marker gene specificity to individual species. For species with more than 100 conspecific genomes, we adjusted the uniqueness criterion to reflect broader presence of the gene. We capped the number of selected marker genes at 200 per species, prioritizing longer sequences for inclusion. Ultimately, we succeeded in identifying marker genes for 1,523 of the 1,524 MRGM species, with one species (MRGM_0393) being an exception. These selected marker genes were compiled into a custom database tailored for MetaPhlAn4, utilizing Bowtie2-build with the ’--large-index’ option to accommodate the database size.

### Generation of simulation datasets for taxonomy profiling method comparison

To facilitate the comparison of taxonomy profiling methods, we generated simulation datasets using CAMISIM ^56^, selecting only genomes for species that are included in both the MGBC ^15^ and the MRGM to prevent bias towards either catalog. We set the sequencing size to 5 Gbp per sample, while keeping all other parameters at their default settings. We constructed synthetic communities comprising 40, 150, and 600 genomes, methodically increasing the genome count to augment the complexity of these communities. To further enhance the complexity of genome selection within these communities, we adopted three strategies: (1) selecting only the representative genome for each species, (2) choosing a genome at random from among conspecific genomes for each species, and (3) selecting five genomes at random from each species. Each genome was assigned an equal abundance to ensure uniform distribution across the samples, thereby generating a diverse array of synthetic communities for subsequent analysis. We organized nine unique setups based on combinations of three community sizes and three genome selection strategies, generating three random samples for each configuration. This resulted in a total of 27 simulation samples. For the methodological comparison, we aggregated results from all simulation samples for each profiling method, allowing for a comprehensive evaluation of their performance.

### Evaluation of taxonomic profiling methods on simulation datasets

We assessed the performance of taxonomic classification across simulation datasets by comparing the effectiveness of different reference databases for mouse gut microbial genomes: the MRGM and the MGBC ^15^. For marker-based taxonomic profiling, MetaPhlAn4^55^ was utilized with a custom-built database derived from MRGM. In contrast, for DNA-based taxonomic profiling, we implemented Kraken2 ^40^ and Bracken ^41^ using custom databases for both MRGM and MGBC. Kraken2 was executed both with and without a confidence threshold set at 0.2. Following the Kraken2 analysis, species-level relative abundances were recalculated using Bracken. To ensure accurate representation, we normalized the relative abundance data based on the genome size of the representative genome that scored highest in genome intactness within each species.

### Evaluation of taxonomic profiling methods on real datasets

We acquired WMS data from research on the diet-associated microbiomes by Manghi et al. ^55^, specifically selecting cross-sectional mouse samples that were not subjected to antibiotic treatments. For our analysis, we excluded the cohort studied by Campbell et al. ^57^ due to its incorporation of both specific pathogen-free (SPF) and germ-free (GF) mice across the different dietary groups.

To evaluate the efficacy of different taxonomy profiling methods, we utilized a random forest classifier to distinguish between mice on high-fat and low-fat diets. Bacterial relative abundances in each sample were normalized using the center log-ratio (CLR) method. This normalized data served as the basis for training random forest classifiers under two scenarios: Cross Study Validation (CSV) and Leave One Dataset Out (LODO). In the CSV scenario, the classifier was trained on one dataset and tested on another, while in the LODO scenario, it was trained on all datasets but one, which was then used for validation. We adopted the random forest algorithm as implemented in metaML ^58^, following the parameters outlined by Thomas et al ^59^. The script for this analysis was obtained from the metaML GitHub repository (https://github.com/SegataLab/metaml).

### Taxonomic comparison between mouse and human gut microbiomes

To compare the gut microbiomes of mice and humans, we utilized data from an updated version of the Human Reference Gut Microbiome (HRGM) ^9^, which includes 4,824 species by NC genomes with a CSS ≤ 0.45 (a manuscript currently under preparation). Taxonomic annotation for the gut bacterial genomes of both mice and humans was carried out using GTDB-Tk version 2.1.1 ^32^. Given the limitations of GTDB-Tk in annotating some species, a direct comparison based solely on the number of shared species might not accurately reflect the true overlap and differences between the mouse and human gut microbiomes. To address this issue and ensure a more accurate taxonomic comparison at species-level, we established the identity of genomes belonging to the same species based on an ANI threshold of ≥ 95%. This methodological approach allows for a more precise evaluation of the taxonomic similarity and distinctions between the microbial communities in the mouse and human gut.

### Functional comparison between mouse and human gut microbiomes

For the functional comparison of gut microbiomes between mice and humans, we utilized pangenomes constructed with panaroo ^53^. When a species was represented by only a single non-redundant genome, that genome was treated as its pangenome. The functional annotation of these pangenomes was carried out using eggNOG-mapper ^44, 45^. To assess the differences in functional capacities between the mouse and human gut microbiomes, we conducted enrichment analyses focusing on KEGG Orthology (KO) terms ^47^, applying Fisher’s exact test alongside Bonferroni correction to adjust for multiple comparisons. Given the potential for low annotation rates of KO terms, we enhanced our analysis by clustering the protein families at 90% similarity using linclust from MMseq2 ^43^. This allowed us to evaluate the functional overlap more comprehensively by comparing the clustered protein families, thereby providing a detailed insight into the distinct and shared functional capabilities within the gut microbiomes of mice and humans.

### Predicting biosynthetic gene clusters (BGCs) in genomes and classifying them into biosynthetic gene cluster families (GCFs)

We identified BGCs in 42,245 non-redundant genomes from MRGM and 155,211 non-redundant genomes from HRGM using antiSMASH v6.1.1 ^60^, resulting in the prediction of 68,655 BGCs in MRGM genomes and 307,995 BGCs in HRGM genomes. To categorize and analyze these BGCs, we clustered the combined total of 376,660 BGCs with BiG-SLiCE ^61^, applying an L2-normalized Euclidean distance threshold of T=0.4 for clustering. To assess the biosynthetic potential differences between mice and humans, we conducted an enrichment analysis focusing on the products of GCFs using Fisher’s exact test with Bonferroni correction. This analysis revealed significant enrichment of specific GCF products in either human or mouse microbiomes, indicating distinct biosynthetic capabilities across the two.

### Availability of data

Raw metagenomic sequencing data are available from the NCBI Sequence Read Archive (SRP335854). Sequences and annotations for bacterial genomes, protein families, 16S regions, a custom Kraken2 database, and a database of species-specific markers are available from https://www.decodebiome.org/MRGM/.

## Results

### MRGM provides high-quality mouse gut bacterial genomes

To develop the MRGC, we integrated pre-assembled genomes from public databases with genomes newly assembled from public WMS data of 1,837 fecal or cecal samples previously unused for de novo genome assembly, using an in-house genome-resolved metagenomic analysis pipeline (**Supplementary Fig. 1a**). Consistent to our previous study ^9^, we found that that ultra-deep sequencing enhances the assembly of MAGs (**Supplementary Fig. 1b**).

Aiming to assemble genomes for novel species with low abundance, we generated WMS data with deep sequencing from 40 newly collected mouse fecal samples. We sequenced these 40 samples to an average depth of 22.4 Gbp, equivalent to approximately 74 million read pairs. Although we did not assemble genomes for novel species from the 40 deep-sequenced samples, we managed to obtain higher quality representative genomes for 116 species. Contigs derived from *de novo* genome assembly of the 1,877 (1,837 public and 40 in-house) WMS datasets (**Supplementary Table 1)** were organized into genome bins using an ensemble binning method, resulting in the generation of MAGs (**Methods**). The completeness and contamination levels of these genome bins were assessed using CheckM2^26^. Consequently, a total of 55,006 MAGs of medium or higher quality were obtained. To further enrich the MRGM, we integrated an additional 60,452 MQ MAGs from the MGBC ^15^ and 202 isolated genomes from the miBC ^29^, as well as 157 single-cell amplified genomes (SAGs) ^30^ (**Supplementary Table 2**).

All collected genomes underwent further quality assessment using the GUNC tool ^28^. GUNC evaluates genomic chimerism based on the homogeneity of contigs and employs an entropy-based metric known as the clade separation score (CSS). We applied a filter for NC genomes with a CSS ≤ 0.45, adhering to the default threshold set by the GUNC software. This process yielded a total of 55,893 genomes for inclusion in the MRGM. We next dereplicated the 55,893 genomes iteratively to 1,524 species clusters (**Supplementary Table 3**). To eliminate redundant genomes within the MRGM catalog, we utilized average-linkage hierarchical clustering with a stringent criterion of 99.9% average nucleotide identity (ANI) and a 0.81 coverage cutoff. This approach effectively refined the MRGM to encompass 42,245 non-redundant, high-quality microbial genomes, distributed across 1,524 species. The completeness and contamination of these genomes are shown in **Supplementary Fig. 1c**. For each species cluster, we identified and selected the highest-quality genomes to serve as representatives.

We then proceeded to taxonomically annotate each representative genome using the Genome Taxonomy Database Toolkit (GTDB-Tk) ^32^. This analysis confirmed that the species diversity within the MRGM catalog is exclusively bacterial, with no archaeal genomes identified among the 1,524 species. The bacterial composition is predominantly characterized by three major groups: Firmicutes_A, with 927 species accounting for 60.8% of the total; Bacteroidota, with 221 species representing 14.5%; and Firmicutes, comprising 167 species or 11.0% of the species diversity (**Fig. 1a**). This composition highlights the significant bacterial diversity captured in the MRGM catalog and underscores its value for gut microbiome research.

**Figure 1.**
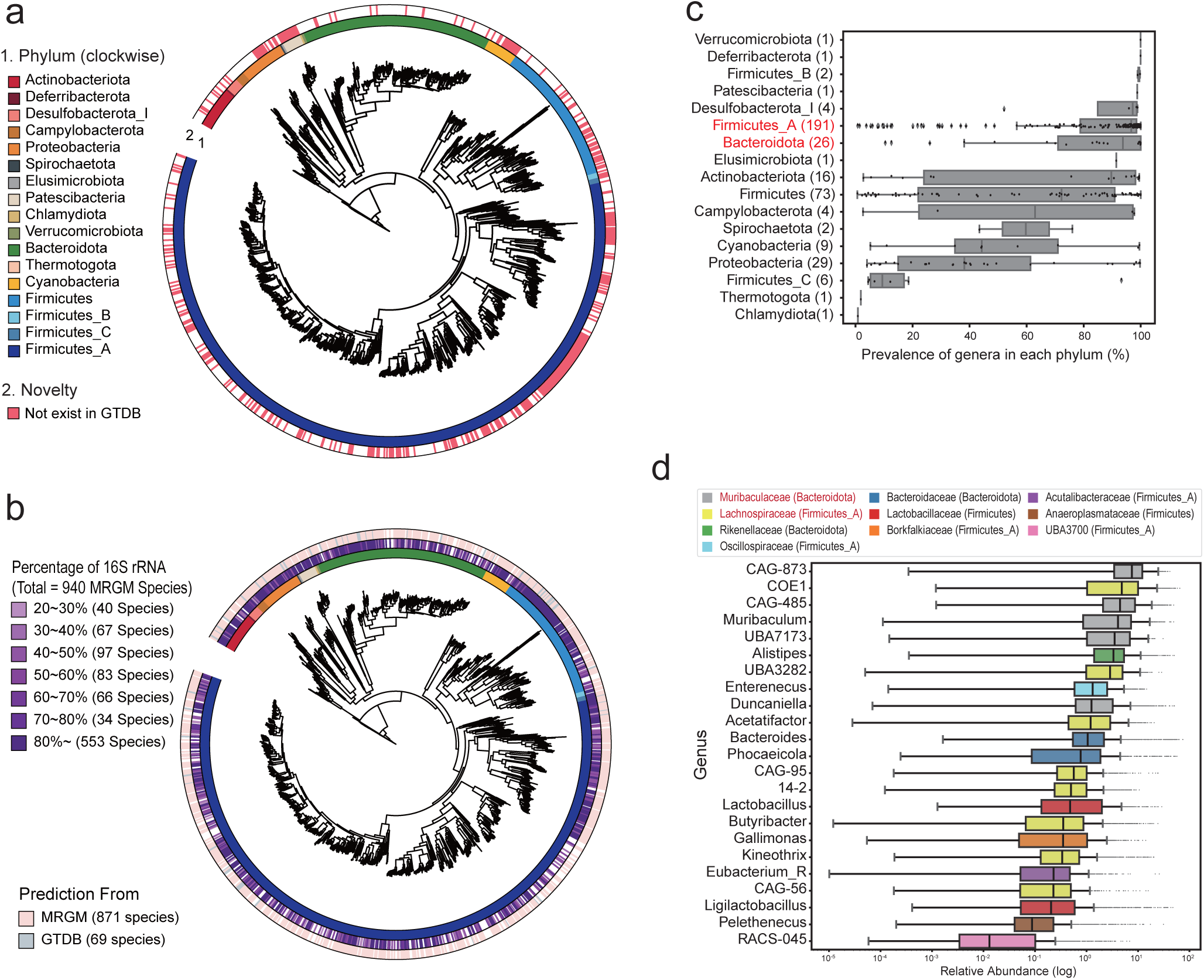
Overview of mouse gut microbial genomes in MRGM. (a) Phylogenetic tree displaying MRGM species diversity; colors indicate phyla and shading denotes novelty (red for MRGM-specific species). (b) Tree showing the proportion of 16S rRNA genes, with color intensity reflecting the percentage presence and source prediction indicated by color codes in the outer ring. (c) Box plot depicting the prevalence of genera within MRGM phyla across 1,843 tested samples, with the number of genera in each phylum noted in parentheses. (d) Box plot of relative abundance for 23 genera that are present in all 1,843 tested samples.

To enable amplicon-based metagenomic analysis, we sought to identify 16S rRNA sequences for each species within the MRGM catalog. For this purpose, we utilized the barrnap software ^39^ to search for 16S rRNA sequences within the genomes. It is well-documented that highly conserved genomic regions, such as 16S rRNA genes, are not typically assembled efficiently from short-read sequencing data due to their conservation across different species ^62, 63^. To improve the likelihood of recovering 16S rRNA sequences, our search extended beyond the representative genomes to include other member genomes within each species cluster. Despite these efforts, 16S rRNA sequences were identified in only 871 of the 1,524 species, representing 46.8% of the total. To address the gaps in 16S rRNA sequence coverage, we then referenced the GTDB r207 genomes corresponding to MRGM species that lacked a 16S rRNA sequence. By analyzing these additional genomes, we succeeded in identifying 16S rRNA sequences for an additional 69 species. Consequently, the MRGM now includes a total of 16,020 16S rRNA sequences for 940 of the 1,524 species, enhancing its utility for amplicon-based metagenomic studies with a coverage of 61.7% (**Fig. 1b**, **Supplementary Table 4**).

Leveraging the MRGM genome catalog, we conducted a survey to determine the prevalence of various genera within mouse gut microbiomes by analyzing 1,843 WMS samples used for MAG construction and had sequencing depths greater than 100Mbp. This investigation aimed to identify the most commonly occurring genera across the datasets, providing insights into the microbial diversity and core constituents of the mouse gut microbiome. Our analysis revealed that two major phyla—Firmicutes_A and Bacteroidota—not only encompass the majority of identified species but also include the most prevalent genera across the mouse gut microbiome (**Fig. 1c**). Specifically, we identified 23 genera that are present in all of the 1,843 mouse gut microbiome datasets. Notably, eight of these universally present genera belong to the *Lachnospiraceae* family within Firmicutes_A, and five are part of the *Muribaculaceae* family within Bacteroidota (**Fig. 1d**). This finding underscores the significance of these families and their associated genera in the mouse gut microbiome, highlighting their potential importance in gut microbiome stability and function.

### Deep learning expands the functional landscape of mouse gut bacterial proteins

In our study, we created a comprehensive catalog of mouse gut bacterial proteins by predicting 132,391,284 coding sequences from 55,893 NC genomes using the Prokka ^42^. Given the prevalence of homologous protein sequences within the microbiome, we aimed to minimize redundancy in our protein catalog. To achieve this, we first clustered protein sequences with 100% sequence identity, excluding exact duplicates, which yielded approximately 11.7 million unique protein sequences. Further clustering at similarity thresholds of 90%, 80%, and 50% resulted in approximately 4.1 million, 3.0 million, and 1.4 million protein sequences, respectively (**Supplementary Table 5**).

Notably, the majority of protein families are species-specific. For example, over 3 million out of the 4.1 million protein families identified at 90% identity are unique to specific species (**Fig. 2a**, indicated by bar colors). This suggests that the speciation of mouse gut bacteria is influenced not only by the presence of different protein contents but also by their sequence variations. Initial functional annotation of these proteins was performed using the eggNOG-mapper tool ^44^. We observed a trend where the rate of functional annotation decreased as protein redundancy decreased (**Fig. 2a**, indicated by lines). This trend likely occurs because proteins with a higher number of homologs are more extensively characterized functionally. Proteins with homologs in different species, for instance, are often more likely to be annotated based on known functions within their orthologous groups ^45^. Conversely, proteins lacking orthologs in other species seldom benefit from homology-based functional annotations.

**Figure 2.**
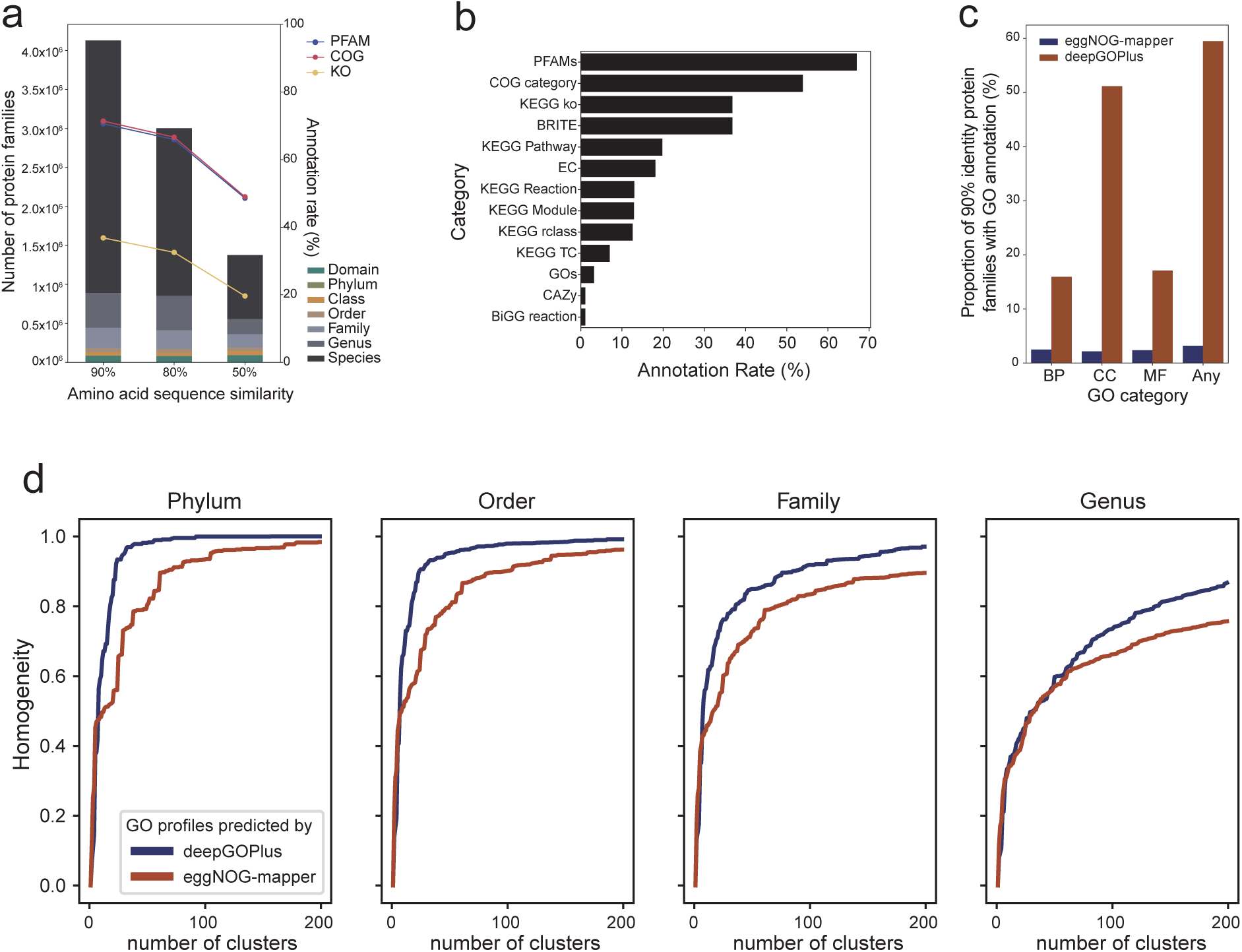
Functional annotations of mouse gut microbial proteins in MRGM. (a) Protein families and annotations at different similarity thresholds, with lines indicating KEGG orthology (KO), protein family (PFAM), and clusters of ortholog group (COG) assignments. Proportion of the lowest taxonomic rank shared for each protein family is indicated by distinct bar colors. Annotation rates are indicated by lines. (b) Annotation rate of 90% identity protein families by various functional categories of eggNOG-mapper. (c) Comparison of annotation rate for each category of GO and any GO terms by eggNOG-mapper and deepGOPlus. (d) Taxonomic homogeneity of clusters by GO annotations by either deepGOPlus or eggNOG-mapper across taxonomic levels (Phylum to Genus).

The eggNOG-mapper facilitates protein annotation using a variety of functional terms, including the protein families (PFAMs) ^46^, Clusters of Orthologous Groups (COGs) ^64^, KEGG Orthology (KO), KEGG pathways ^47^, Gene Ontology (GO) ^48^, enzyme commission (EC) numbers ^50^ and Carbohydrate-Active enZymes (CAZy) ^49^. We noted acceptable annotation rates for PFAMs and COGs for protein families at a 90% similarity threshold. Nevertheless, PFAM terms and COG categories provide only broad functional categorizations, making it challenging to deduce the specific functional roles of proteins in bacterial physiology. GO annotations are a cornerstone in the study of gene functions, offering a framework that is extensively utilized across biological research. This system of annotation is particularly valuable for exploring the functional roles of proteins and for generating novel hypotheses concerning mouse gut microbial proteins. Yet, we were only able to annotate a meager 3.2% of protein families using GO terms (**Fig. 2b**). To enhance the GO functional annotation of mouse gut microbial proteins, we utilized the established deep learning method deepGOPlus ^51^, which leverages a convolutional neural network (CNN) to learn representations capable of predicting GO terms from protein sequences. Noticing that some predicted eukaryotic cellular functions were not applicable to bacterial proteins, we filtered out eukaryote-specific GO terms to ensure only bacterial protein functions were considered. Impressively, this deep learning method significantly increased the GO annotation rate for the 90% identity protein family from 3.2% with orthology to around 60%, achieving more than 18-fold increase (**Fig. 2c**). This substantial improvement demonstrates that deep learning can surmount the constraints of homology-based functional annotations, providing a much broader and detailed functional landscape of mouse gut microbial proteins.

To assess the quality of functional annotations provided by deepGOPlus, we analyzed 1,524 representative genomes from each species based on GO terms obtained from both deepGOPlus and eggNOG-mapper. We then clustered these genomes according to their GO profile distances, applying various thresholds for agglomerative clustering (**Methods**). The homogeneity of taxonomic classes within clusters was determined by the consistency of genus labels among the genomes within each cluster, extending this approach to higher taxonomic levels as well. Our findings revealed a higher degree of homogeneity within clusters formed based on GO term profiles from deepGOPlus compared to those from eggNOG-mapper (**Fig. 2d**). Given the assumption that species taxonomically closer are likely to share more similar functional contents, these results highlight the superiority of deepGOPlus in providing functional annotations for the encoded proteins of gut microbial genomes.

### MRGM significantly broadens taxonomic diversity of the mouse gut microbiota

The MRGM catalog exclusively comprises microbial species with NC genomes because the completeness of a genome significantly influences downstream functional analysis, often leading to underestimation of its functional capacity ^65^. Unlike most previously published catalogs of mouse gut microbial genomes that include species with MQ genomes ^13, 14, 16^, MGBC, a recently published catalog ^15^, stands out by containing the largest number of NC genomes—26,640 genomes across 1,094 species. This distinction makes MGBC a suitable benchmark for evaluating improvements in the MRGM over earlier catalogs.

Our comparison revealed that 1,051 species are common to both the MRGM and the MGBC. Additionally, MRGM encompasses 473 species not found in MGBC, whereas MGBC contains only 39 species not included in MRGM. Notably, of the 39 species unique to MGBC, 38 failed to meet the NC genome criteria set by CheckM2 ^26^ and GUNC ^28^, (**Supplementary Table 6**). The 473 species unique to MRGM are distributed across the entire phylogenetic tree, highlighting the taxonomic breadth of the novel genomes (**Fig. 3a**). Interestingly, a significant portion (34% or 161 of the 473) of MRGM-exclusive species showed a strong association with specific lineages—including the phylum Patescibacteria, family *Gastranaerophilaceae*, phyla Firmicutes_B and Firmicutes_C, family *UBA1242*, and order TANB77—that almost exclusively exist in MRGM (**Fig. 3b**). This pattern suggests that the genomes unique to MRGM, associated with these lineages, benefit from the advanced genome quality assessment methodology employed for MRGM, which was not used for previously published catalogs including MGBC.

**Figure 3.**
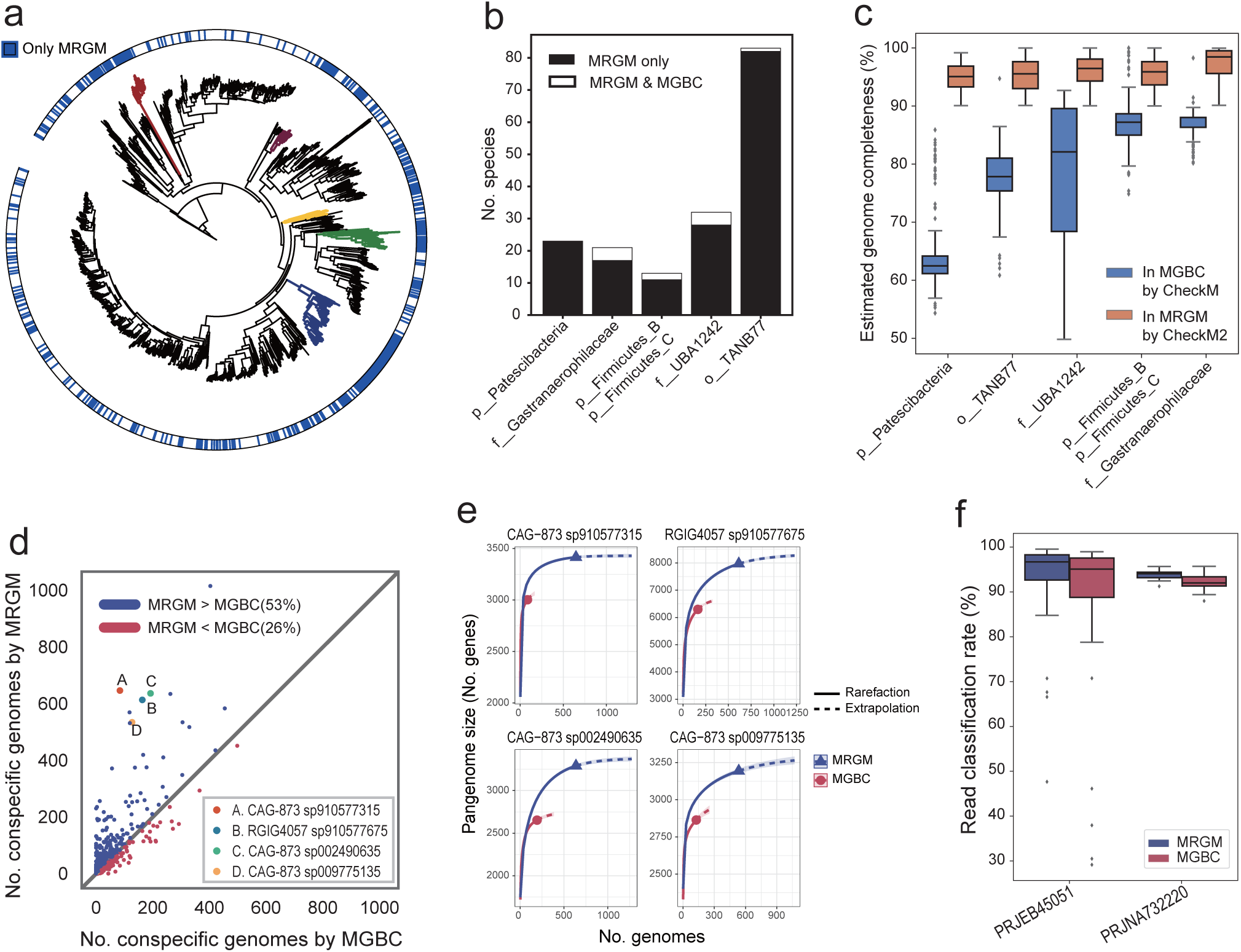
Enhanced taxonomic diversity of mouse gut microbiota. (a) A phylogenetic tree delineating MRGM-exclusive and shared species with MGBC indicated in the outer ring. Some lineages predominantly unique to MRGM are highlighted in different colors. (b) Bar graphs showing the number of species unique to MRGM and shared with MGBC across the highlighted lineages. (c) Box plots comparing estimated completeness of genomes for each lineage in MGBC and MRGM using CheckM and CheckM2, respectively. (d) A scatter plot comparing the number of conspecific genomes for common species in MRGM versus MGBC. The four species with significantly increased number of conspecific genomes in MRGM are indicated as A, B, C, and D. (e) Rarefaction and extrapolation curves for pangenome size of four selected species within MRGM and MGBC. (f) Box plots illustrating taxonomic classification rates for metagenomic sequencing reads from two datasets (PRJEB45051, PRJNA732220) by MRGM and MGBC.

For a long time, CheckM has been the standard tool for evaluating genome quality, specifically in terms of completeness and contamination ^27^. Despite its reliability for a broad range of genomes, CheckM, which relies on lineage-specific single-copy marker genes, encounters difficulties accurately estimating the completeness of genomes belonging to novel or under-represented lineages ^66^. This limitation stems from the absence of specific marker genes in such lineages, leading to potential underestimations of genome completeness. Addressing this challenge, an improved version, CheckM2, has been developed, incorporating machine learning techniques to refine its assessment process ^26^. CheckM2 allows for a more accurate evaluation of genome completeness across a wider array of lineages, including those previously under-sampled or entirely novel to existing catalogs. By employing CheckM2 in our bin refinement and genome quality assessment processes, we were able to include high-quality genomes from lineages not represented in prior catalogs of high-quality microbial genomes for the mouse gut microbiome. Where CheckM previously underestimated the completeness of MAGs from novel lineages, CheckM2 has facilitated the incorporation of numerous genomes from these previously overlooked lineages (**Fig. 3c**). This advancement has markedly improved the MRGM catalog over MGBC, expanding taxonomic diversity and the scope of mouse gut microbial genomes it encompasses.

Even within a single species, different strains can display diverse functional capacities ^67–70^. Consequently, possessing a larger collection of conspecific genomes for each species is advantageous. We assessed the genome counts for each of the 1,051 species shared between the MRGM and the MGBC, discovering that 53% of these species feature a greater number of genomes in MRGM compared to MGBC (**Fig. 3d**). This increase in conspecific genomes is anticipated to expand the pangenome size for each species. Utilizing panaroo ^53^, we generated pangenomes for each species and observed significant expansions in the pangenomes of many species, showcasing four prominent examples (*CAG-873 sp910577315*, *RGIG4057 sp910577675*, *CAG-873 sp002490635*, and *CAG-873 sp009775135*) where MRGM demonstrated a marked advantage over MGBC (**Fig. 3e**). Through rarefaction analysis, we verified that the pangenome sizes of these species approached saturation within MRGM, an outcome not observed with MGBC.

The accuracy of DNA-based taxonomic profiling of metagenomic data is highly dependent on the quality and scope of the reference genome database ^71^. Given that the MRGM catalog is more comprehensive than previous databases, we anticipated an enhancement in the taxonomic classification of metagenomic sequencing reads. To evaluate this, we employed Kraken2 ^40^ with custom databases derived from the concatenated genomes for each species within MGBC and MRGM. Our assessment used 64 WMS samples by paired-end sequencing from PRJEB45051 and 16 WMS samples by single-end sequencing from PRJNA732220, none of which were involved in the development of either genome catalog. We observed a significant enhancement in sequence read classification rates using MRGM compared to MGBC (**Fig. 3f**, *P* < 0.001, Wilcoxon signed-rank test), potentially leading to more accurate taxonomic profiling of mouse gut metagenomes.

### MRGM enables accurate taxonomic profiling with both DNA- and marker-based methods

For analyzing metagenomic sequencing data, the DNA-based Kraken2 ^40^, often used in conjunction with Bracken ^41^, and the marker-based MetaPhlAn ^72^ are commonly utilized ^73^. Kraken2 offers ease of integration with genome catalogs due to its straightforward process for creating custom databases of DNA sequences with their taxonomic information. In contrast, MetaPhlAn demands more effort to derive species-specific markers from the genome catalog, which may partially explain why MGBC provided a custom database for Kraken2 but not for MetaPhlAn. To enhance the versatility of MRGM in taxonomic profiling, we have created both a custom Kraken2 database and a database of species-specific marker genes, utilizing sequences from the MRGM genomes (**Methods**).

To assess the species-level taxonomic profiling capabilities of MRGM compared to MGBC, we created simulated metagenomic sequence reads from genomes common to both MGBC and MRGM using CAMISIM ^56^ (**Methods**). We then applied different taxonomic profiling pipelines: MGBC or MRGM with Kraken2, supplemented by Bracken for refined estimation of species abundance (denoted as MGBC + Kraken2 and MRGM + Kraken2, respectively), and MRGM with MetaPhlAn (MRGM + MetaPhlAn). Kraken2-based analyses must consider additional factors, such as the tendency for species with larger genomes to yield more aligned reads, thus appearing more abundant ^74^. Additionally, the impact of ambiguously assigned reads by Kraken2 significantly affects relative abundance of species ^75^. Therefore, our analysis also examines the effects of genome size normalization and the implementation of a Kraken confidence score threshold (excluding reads with a confidence score below 0.2).

We observed that while all tested pipelines achieved perfect recall (zero false negatives), the MRGM + MetaPhlAn and MRGM + Kraken2, when enhanced with genome size normalization and a confidence score cutoff, exhibited superior precision and Bray-Curtis similarity (**Fig. 4a**). Both MRGM and MGBC performed poorly in species-level taxonomic profiling when using Kraken2 without genome size normalization or confidence score cutoff. These findings indicate MRGM has the capacity to provide more accurate taxonomic profiling of metagenomic data than MGBC through both DNA- and marker-based approaches.

**Figure 4.**
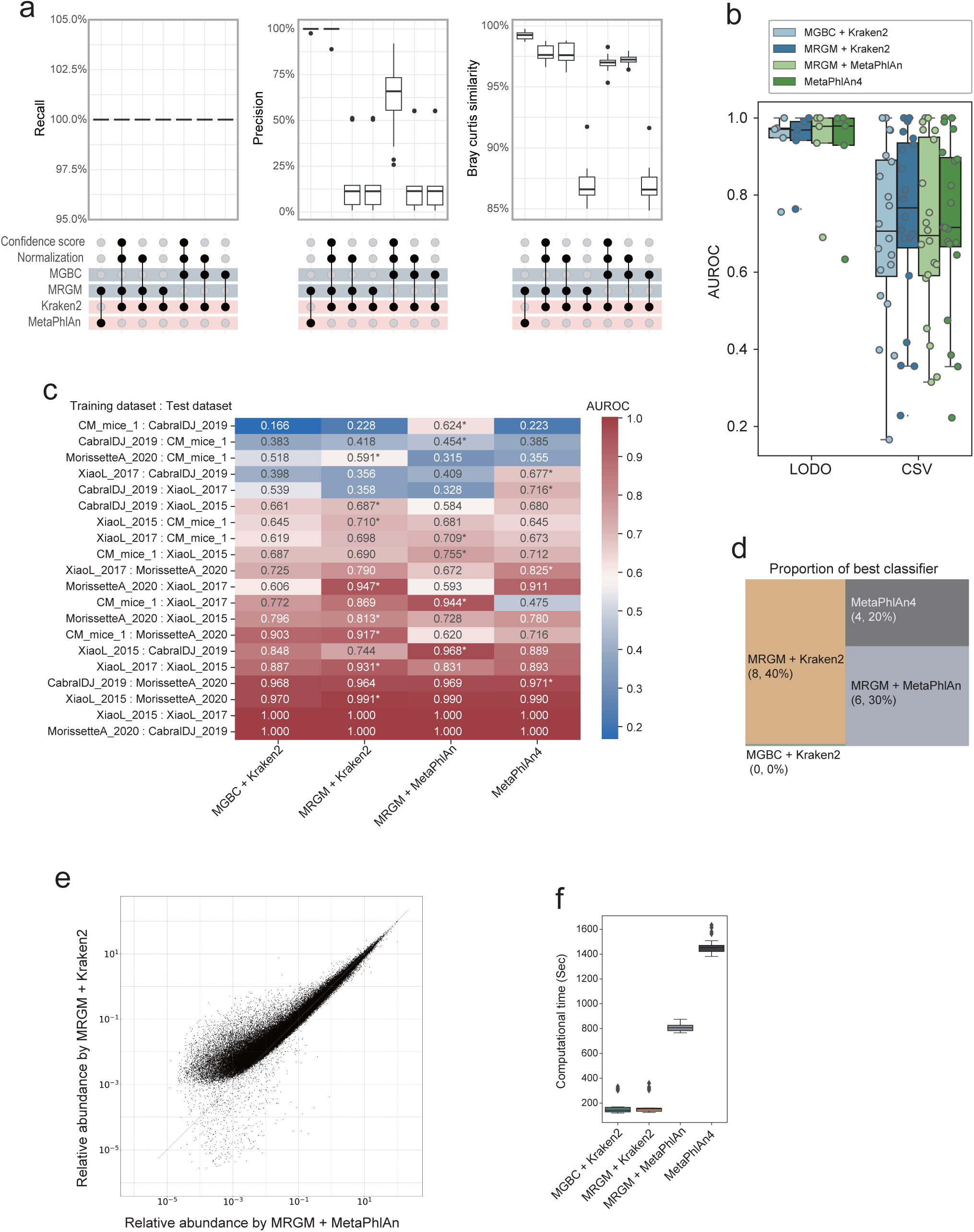
Assessment of taxonomic profiling with MRGM. (a) Box plots illustrating recall, precision, and Bray-Curtis similarity metrics for classification of metagenomic sequencing reads of 27 simulation datasets into source species using MRGM and MGBC genomes combined with Kraken2 and MetaPhlAn methods under various conditions including confidence score cutoff and genome size normalization. (b) Box plots depicting the Area Under the Receiver Operating Characteristic (AUROC) scores of the random forest classifiers which predicts high-fat and low-fat diet mouse gut microbiota of real data in two settings, Leave One Dataset Out (LODO) and Cross Study Validation (CSV). (c) A heatmap displays the AUROC values for random forest classifiers for predicting mouse diet across different training and test dataset combinations, showcasing the relative accuracy of each method. The x-axis labels the classification methods, while the y-axis details the datasets used for training and testing. An asterisk marks the best-performing method for each dataset combination. (d) A graph representing the proportion of times each classifier was determined to be the best performer in distinguishing mice fed with a high-fat diet and a low-fat diet in real metagenomic sequencing datasets. (e) A scatter plot comparing the relative abundance of taxa as determined by MRGM with Kraken2 versus MRGM with MetaPhlAn. (f) Box plots comparing computational times (with four CPU cores) required for metagenomic classification by each method using simulation dataset.

Furthermore, we evaluated taxonomic profiling capabilities of MRGM and MGBC using real metagenomic sequencing data from diet-related microbiome studies across multiple cohorts (**Supplementary Table 7**). For the assessment with real data, we also included MetaPhlAn4, utilizing its own species-specific marker database, ChocoPhlAn ^76^, which is not tailored to the mouse gut microbiome. Taxonomic profiling performance was evaluated by distinguishing microbiomes associated with high-fat and low-fat diets through species abundance profiles using random forest classifiers in two approaches: Cross Study Validation (CSV), where the model is trained on one dataset and validated on a distinct dataset not involved in training, and Leave One Dataset Out (LODO), which trains the model on all datasets except one reserved for validation. While performance was high across all pipelines with LODO, MRGM + Kraken2 outperformed MGBC + Kraken2, MRGM + MetaPhlAn, and MetaPhlAn4 in CSV, albeit without statistical significance (**Fig. 4b**). MRGM consistently showed the best performance (**Fig. 4c**), leading in 8 and 6 out of 20 CSV scenarios with MRGM + Kraken2 and MRGM + MetaPhlAn, respectively (40% and 30%). In contrast, MGBC + Kraken2 never achieved the best performance (**Fig. 4d**). Notably, in MetaPhlAn-based profiling, MRGM custom species-marker database tailored to the mouse gut microbiota surpassed the performance of the MetaPhlAn4 database, which contains markers from a significantly larger pool of genomes across various habitats, including but not limited to the mouse gut. This underscores the effectiveness of genome databases for specific ecosystems in taxonomic profiling.

We posited that the MRGM genome catalog performance varies between DNA- and marker-based methods due to differences in their sensitivity towards detecting taxa of low abundance. Theoretically, methods that utilize the entire genome for taxonomic detection are more likely to identify taxa, particularly those at low abundance, than methods relying solely on specific marker sequences. To test this hypothesis, we analyzed the relationship between the estimated relative abundance of mouse gut bacterial species using MRGM reference genomes via MetaPhlAn and Kraken2. As expected, Kraken2 revealed a higher estimated relative abundance for many species compared to MetaPhlAn, notably among species of lower abundance (**Fig. 4e**).

As sequencing depth and sample size increase, the demand for computational resources for taxonomic profiling becomes a critical consideration. Our findings indicate that DNA-based profiling methods, such as Kraken2, require significantly less computational time compared to marker-based approaches like MetaPhlAn, which are more time-intensive (**Fig. 4f**).

In summary, we developed custom databases of MRGM genomes tailored for both DNA-based and marker-based methods of taxonomic profiling. Our findings reveal that utilizing MRGM with both approaches surpasses the accuracy of prior genome catalogs, as evidenced by both simulated and real data analyses.

### Comparative analysis of genome catalogs unveils dietary-related taxonomic differences in gut microbiota between mice and humans

Previous catalogs of mouse gut bacterial genomes indicated that around 10% of mouse gut bacterial species are shared with the human gut. Using an enhanced version of the Human Reference Gut Microbiome (HRGM) ^9^, which comprises only high-quality genomes for 4,824 species, we investigated overlap of common gut microbial species between mice and humans. In line with previous research, our analysis with the MRGM revealed that merely 11.8% (180 out of 1,524) of species are shared with those in the HRGM (**Supplementary Table 8**). This limited species-level taxonomic overlap between human and mouse gut microbiota may highlight their distinct evolutionary adaptations to their respective host gut environments. Consequently, we explored the differences across various taxonomic levels between human and mouse gut microbiota.

We noted a higher proportion of genera within the mouse gut microbiota that are also present in the human gut microbiota (249 out of 367, 67.8%) (**Fig. 5a**). However, when considering the mouse gut microbiota as a model for the human gut microbiome, it is crucial to account for both taxa shared with humans and those taxa that are absent in the mouse gut. We observed a significantly lower proportion of shared genera in the human gut microbiota appearing in the mouse gut microbiota (249 out of 1,071, 23.2%) (**Fig. 5b**). For instance, while all species of mouse gut bacteria within the Verrucomicrobiota phylum, including the mucin-degrading *Akkermansia muciniphila* ^77^, are found in the human gut microbiota, this phylum in humans encompasses an additional 16 genera. Furthermore, seven phyla present in the human gut microbiota lack representatives in the mouse gut microbiota. Notably, the Fusobacteriota phylum, which includes *Fusobacterium nucleatum*—a species linked to human colorectal cancer ^78^—is absent in the mouse gut microbiota. These findings suggest that the human gut microbiota have developed into a broader array of lineages, resulting in an expanded taxonomic diversity compared to mouse gut microbiota, likely reflecting the more varied human diet.

**Figure 5.**
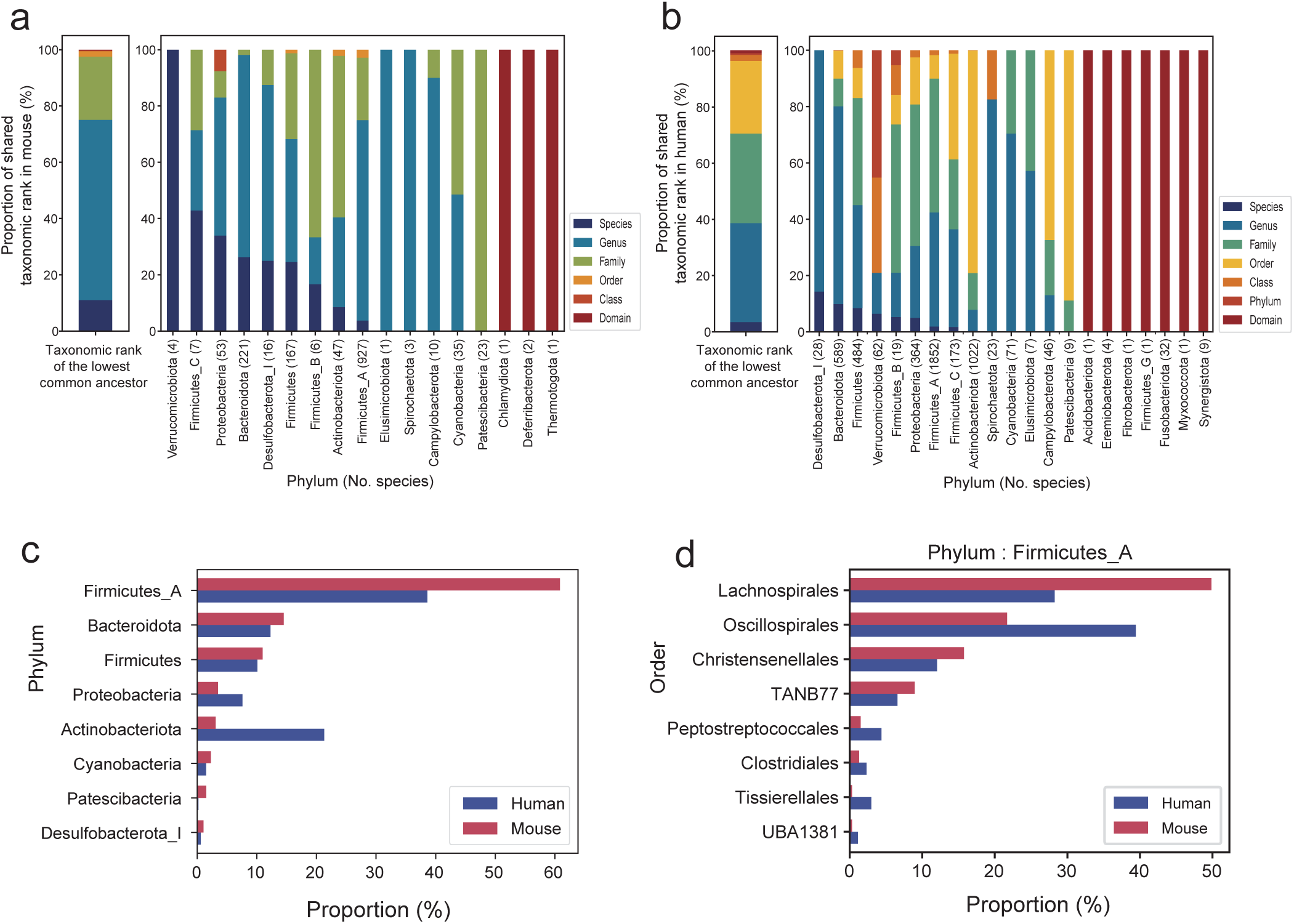
Taxonomic divergence between human and mouse gut microbiota. (a-b) Stacked bar graphs showing distribution of gut microbiota phyla across mouse (a) and human (b), color-coded for the lowest common taxonomic ranks. (c) Bar graphs showing phylum prevalence in human and mouse gut microbiota (d) Bar graphs showing order prevalence of Firmicutes_A phylum in human and mouse gut microbiota.

We noted distinct compositions of gut microbiota phyla between mice and humans (**Fig. 5c**). In mice, the predominant phylum in the MRGM, Firmicutes_A, represents 60.8% of the gut bacteria, in contrast to 38.4% in humans according to the HRGM. This discrepancy could be partially attributed to the greater presence of Lachnospirales within mice, the most populous order of Firmicutes_A (**Fig. 5d**), known for its critical role in producing short-chain fatty acids (SCFAs) from dietary fibers ^79^. This difference likely reflects the fiber-rich natural diet of mice, which confers a selective advantage to SCFA-producing bacteria in their gut environment. Conversely, Actinobacteriota ranks as the second largest phylum in the HRGM, accounting for 21.2% of human gut bacteria but only 3.1% in the MRGM (**Fig. 5c**). A genus within Actinobacteriota, *Bifidobacterium*, significantly influences host health through interactions with immune system ^80^. Our analysis revealed 22 species of *Bifidobacterium* in the human gut microbiota compared to merely three species in the mouse gut. The preferential nourishment of beneficial gut bacteria, including *Bifidobacterium*, by human milk oligosaccharides (HMOs) ^81^ suggests a coevolutionary relationship that might explain the notably higher proportion of *Bifidobacterium* in the human gut microbiota relative to mice. Overall, our taxonomic comparison between human and mouse gut microbiota underscores the influence of host diet in determining the taxonomic makeup of gut microbiota.

### Protein catalog comparison highlights dietary-driven functional divergence in gut microbiota between mice and humans

Previous research has reported a significant overlap in protein functions between mouse and human gut microbiota. In our study, we validated the functional potential similarity between MRGM and HRGM through KO, GO, and CAZy annotations (88.9%, 90.3%, and 96.3% overlap, respectively) (**Fig. 6a**). Nevertheless, these findings pertain solely to annotated proteins, constituting just 40.2%, 4.2%, and 1.6% of the total for KO, GO, and CAZy annotations, respectively. To delve deeper into the functional similarities between human and mouse gut microbiota beyond these annotated proteins, we explored the overlap of protein families based on 90% identity in amino acid sequences. Our analysis revealed that a mere 16.8% of MRGM protein families (562,205 out of 3,336,320) are shared with HRGM protein families (**Fig. 6b**). This suggests that the previously reported functional overlap between mouse and human gut microbiota may be overestimated due to a focus on proteins with known functions. For the uncharacterized ’functional dark matter’ of gut microbiota, there appears to be a significant divergence between mice and humans.

**Figure 6.**
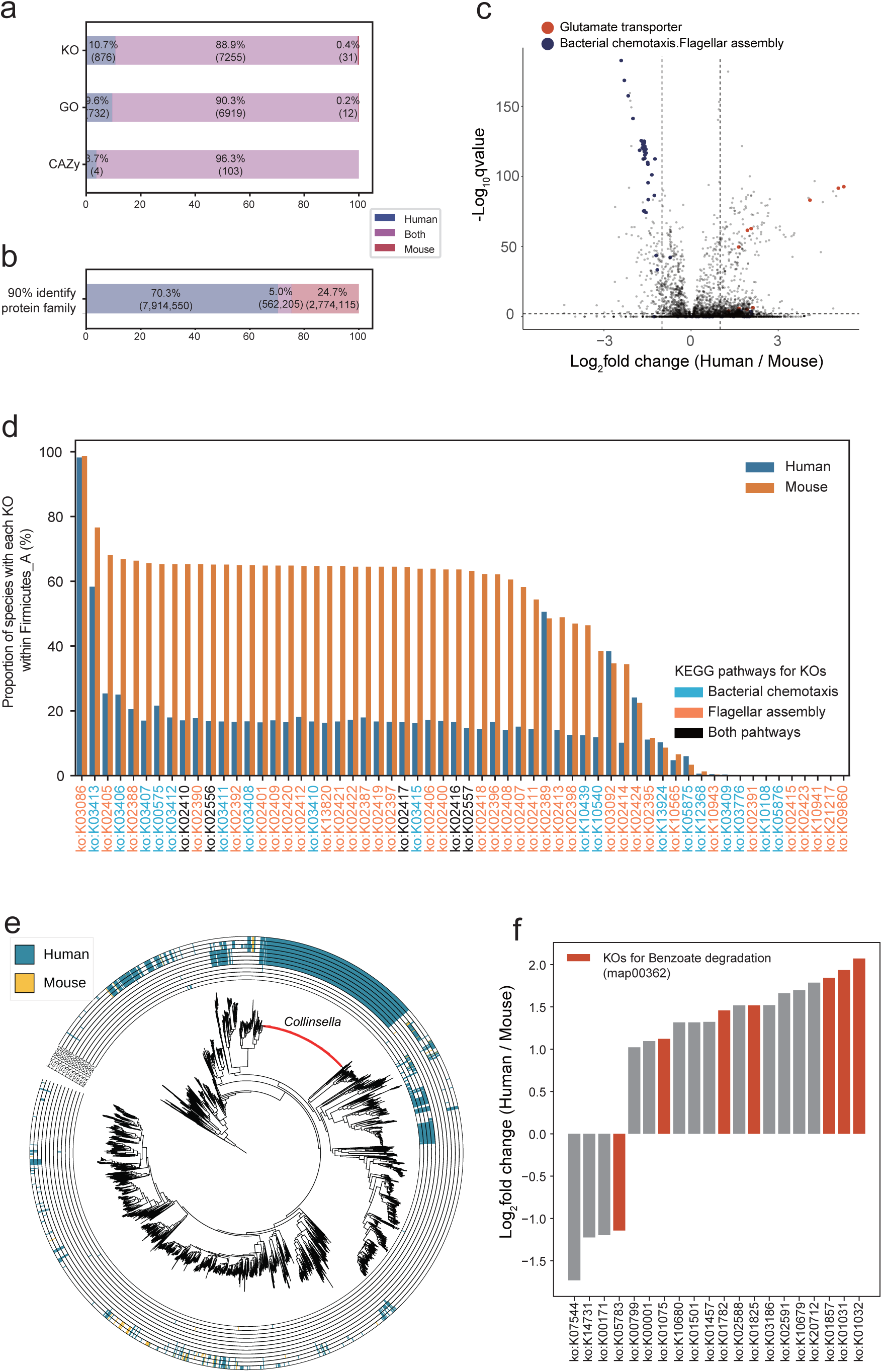
Functional divergence between human and mouse gut microbiota. (a) Stacked bar charts showing the percentage of KEGG Orthology (KO), Gene Ontology (GO), and Carbohydrate activation enzyme (CAZy) annotations unique to human, mouse, or shared by both. (b) Proportion of protein families at 90% amino acid identity, indicating the percentage shared between human and mouse gut microbiota. (c) A volcano plot highlighting differentially enriched KOs between human and mouse microbiota, with points representing KOs related to glutamate transporter, bacterial chemotaxis and flagellar assembly. (d) Bar charts depicting the proportion of species with KOs for pathways associated with bacterial chemotaxis (map02030) and flagellar assembly (map02040) in Firmicutes_A phylum, comparing human and mouse microbiota. (e) A phylogenetic tree for combined set of mouse and human gut microbial genomes with marked species with KOs for glutamate transporter. Species for *Collinsella* genus are indicated by red branches. (f) Bar graphs showing the log-fold change of KOs for xenobiotic degradation between human and mouse gut microbiota, indicating differential enrichment. KOs related to benzoate degradation are marked by red.

Among the functions shared by mouse and human gut microbiota, there could be distinct levels of enrichment across different functional categories for either group. To investigate this aspect, we examined the fold changes in the proportion of species harboring each KO between MRGM and HRGM, identifying several KOs that were differentially enriched between the two (**Fig. 6c, Supplementary Table 9**). Notably, a higher proportion of bacterial species in the MRGM possessed KOs associated with pathways involved in ’bacterial chemotaxis’ (map02030) and ’flagellar assembly’ (map02040). This differential enrichment was especially marked within the phylum Firmicutes_A (**Fig. 6d**), where about 60% of MRGM species exhibited numerous KOs related to chemotaxis and flagellar assembly, compared to only about 20% within HRGM Firmicutes_A. Recent functional comparisons in the human gut microbiota among omnivorous, vegan, and vegetarian volunteers have linked KOs for ’chemotaxis’ and ’flagellar assembly’ with vegan and vegetarian diets ^82^. These insights, combined with our results, imply that the enhanced presence of gut bacterial protein functions related to chemotaxis and flagella assembly in mice correlates with their fiber-rich diet, in contrast to the typical human diet.

We also observed numerous KOs underrepresented in the mouse gut microbiota relative to humans. Specifically, bacterial species associated with ’glutamate transporter’ KOs were less prevalent in MRGM than in HRGM (**Fig. 6c**). Notably, analysis of glutamate transporter KOs (**Supplementary Table 10**) highlighted that the *Collinsella* genus, exclusive to humans, significantly harbors glutamate transporter proteins (**Fig. 6e**). Given that many processed foods consumed by humans contain monosodium glutamate (MSG) as a common additive, humans likely ingest more glutamate in the form of MSG compared to mice. A study assessing the impact of long-term MSG consumption noted an increase in the relative abundance of *Collinsella* with higher MSG intake ^83^. This suggests that glutamate transporters within the *Collinsella* genus afford them a competitive edge in the human gut, where glutamates are abundantly available as MSG, thus underscoring the diet-driven functional and taxonomic divergence between mice and humans.

Humans encounter a broader spectrum of xenobiotics, including antibiotics, diverse drugs, pollutants, food additives, and various chemicals, more so than mice, despite laboratory mice also being exposed to antibiotics. Consequently, we speculated that KOs associated with the degradation and metabolism of xenobiotics might be more prevalent in the human gut microbiota. Our analysis of genes involved in xenobiotic degradation and metabolism revealed a significant enrichment of genes related to benzoate degradation in HRGM (**Fig. 6f**). Sodium benzoate, a common benzoate compound, is extensively used as a food preservative for its antibacterial properties. Frequent exposure of the human gut microbiota to such preservatives may have led to the development of resistance and the widespread acquisition of capabilities to metabolize them.

### MRGM reveals functional divergence in secondary metabolite production between human and mouse gut microbiota

Bacteria synthesize secondary metabolites, bioactive compounds pivotal for both inter-microbial and host-microbe interactions ^84^. These metabolites produced by gut microbiota exhibit significant therapeutic potential including antimicrobial, anticancer, and immunosuppressive properties ^85^. The capacity to generate secondary metabolites is a key functional trait of microbiota, underscoring the value of examining secondary metabolite production across human and mouse gut microbiota to understand their functional differences. To this end, biosynthetic gene clusters (BGCs), which harbor the genetic blueprints for secondary metabolite production, were analyzed. Using antiSMASH ^60^, we identified 68,655 BGCs in the MRGM and 307,995 BGCs in the HRGM genomes. To streamline our analysis and minimize BGC redundancy, we aggregated 376,660 BGCs into 5,883 biosynthetic gene cluster families (GCFs) using BiG-SLiCE ^61^, with 2,040 GCFs associated with the mouse and 5,132 GCFs with the human gut microbiota. This comprehensive survey of BGCs provides a foundational step towards elucidating the secondary metabolite landscape and its functional implications in gut microbiota.

We analyzed the distribution of GCFs across the 1,524 species within the MRGM (**Fig. 7a**). A significant portion of GCFs was identified within the Firmicutes_A phylum, a predominant phylum in the mouse gut microbiota, marking it as a key producer of secondary metabolites. Upon assessing the highest common taxonomic rank associated with each GCF, approximately 60% were unique to specific species, and around 70% were exclusive to particular genera, suggesting that the distribution of GCFs in the mouse gut microbiota is highly specific at the species or genus level (**Fig. 7b**). This taxonomic specificity indicates that GCFs, which categorize secondary metabolite production capabilities, likely signify the unique roles in inter-microbial and host-microbe interactions played by each species or genus. Further analysis of the genera rich in GCFs revealed that the top ten genera with the most GCFs are members of the *Lachnospiraceae* family within the Firmicutes_A phylum (**Fig. 7c**), highlighting their significant contribution to the secondary metabolite landscape of the mouse gut microbiota.

**Figure 7.**
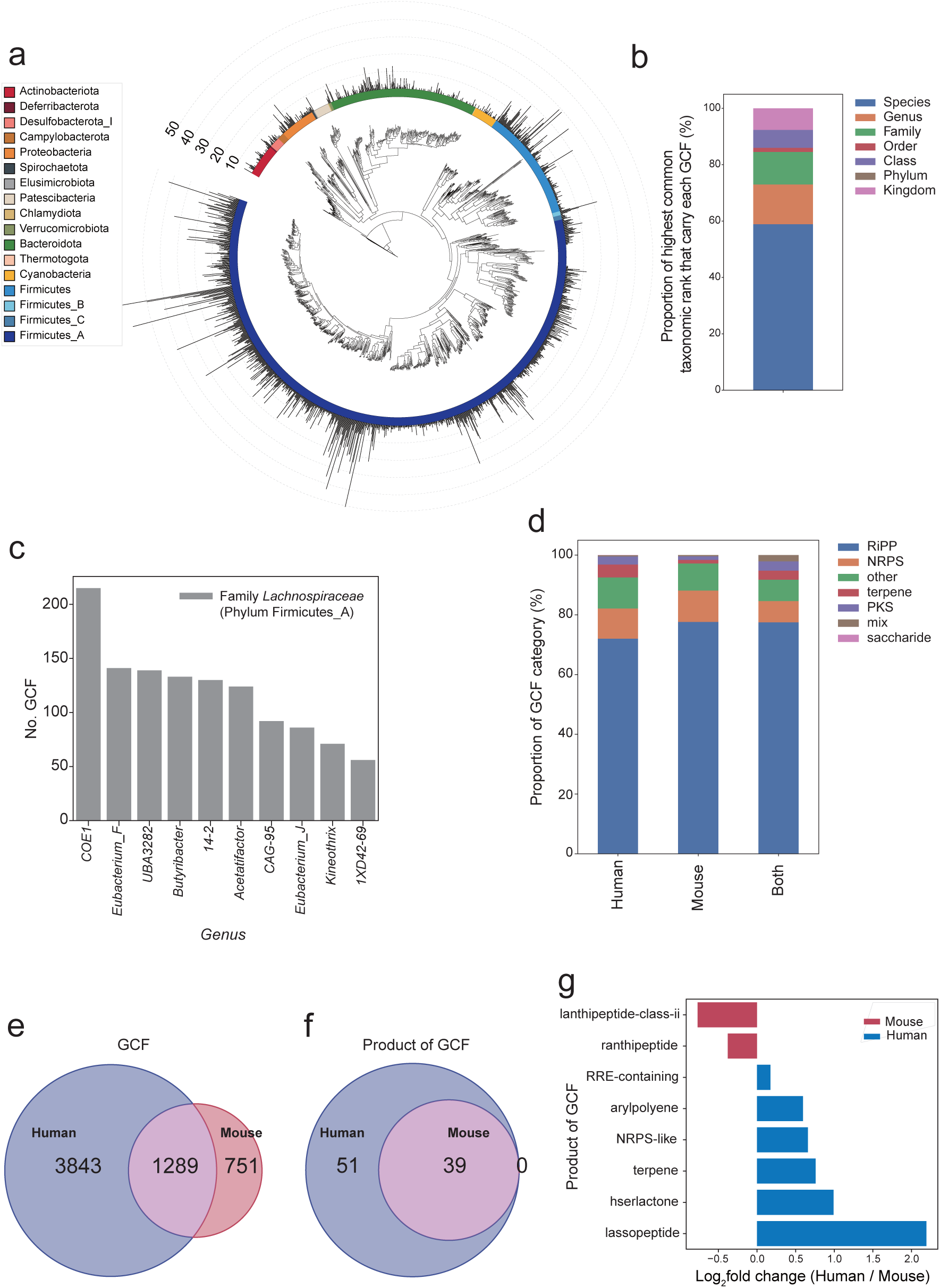
Comparison of secondary metabolite potentials between human and mouse gut microbiota. (a) A phylogenetic tree illustrating the distribution of biosynthetic gene cluster family (GCF) across phyla in the mouse gut microbiota, with a color-coded ring indicating phylum classification. (b) A stacked bar chart displaying proportion of highest common taxonomic rank that carry each GCF in mouse gut microbiota. (c) Bar graphs showing the number of GCFs associated with the top ten genera of mouse gut microbiota, all of which belong to *Lachnospiraceae* family of Firmicutes_A phylum. (d) Stacked bar charts of the proportion of GCF classes between human and mouse microbiota, with a breakdown of GCF classes (e-f) Venn diagram depicting the overlap of GCFs (e) and GCF products (f) between human and mouse gut microbiota. (g) Bar graphs showing the log2 fold change of GCF products between human and mouse gut microbiota, indicating differential enrichment.

In our comparative analysis of secondary metabolite potential between mouse and human gut microbiota, we noted similar distributions of GCF categories across both microbiomes (**Fig. 7d**). This parallel extends to the shared GCF categories, demonstrating that mouse and human gut microbiota have comparable capabilities in broad GCF categories. Despite this overarching similarity, the specific commonality of individual GCFs is more limited, with only 63.2% (1,289 of 2,040) of GCFs shared within the mouse gut microbiota and 25.1% (1,289 of 5,132) within the human gut microbiota common to both (**Fig. 7e**). Intriguingly, although all GCF product categories present in the mouse microbiota were also observed in humans, the human microbiota displayed over twice the variety of GCF product categories than that found in mice (**Fig. 7f**). This discrepancy suggests a fundamental similarity in the production of secondary metabolites, yet with a divergence in the regulatory mechanisms governing these processes between the two microbiomes. Further analysis revealed differential enrichment of specific GCF products: lassopeptides were more abundant in the human gut microbiome, whereas lanthipeptides class ii were predominantly found in the mouse gut microbiome (**Fig. 7g**). Through a systematic comparison of MRGM and HRGM, we uncovered a functional divergence in the production of secondary metabolites between the mouse and human gut microbiomes.

## Discussion

In this study, we present the MRGM, an enhanced catalog of high-quality mouse gut microbial genomes. MRGM includes 1,524 bacterial species, which incorporates 473 novel species, thereby expanding the taxonomic diversity by over 40% compared to the previously most comprehensive catalog, MGBC. Notably, we uncovered that 34% (161 of 473) of these novel species belong to lineages that were hitherto underrepresented, indicating the critical role of enhanced genome quality assessment methods, such as the improved CheckM protocol for MAGs, in capturing these elusive taxa.

MRGM not only broadens the taxonomic diversity but also significantly enriches the functional landscape of the mouse gut microbiota. By employing a deep learning approach, we increased the GO annotation rate from 3.2% with orthology to 60%, achieving more than 18-fold increase.

Custom databases leveraging MRGM genomes have outperformed previous catalogs in accuracy for both DNA- and marker-based taxonomic profiling. Notably, our findings indicate that DNA-based methods, when enhanced with genome size normalization and confidence score filtering, excel in analyzing real data, especially in identifying low-abundance taxa. Furthermore, in marker-based profiling, MRGM surpasses MetaPhlAn4— despite the latter’s use of a significantly larger pool of marker genes from various microbial habitats beyond the mouse intestine—likely due to a reduction in false positives. This highlights the importance of specialized, community-specific genome catalogs for precise taxonomic profiling in whole metagenome sequencing data.

In our comparative analysis, we delve into the taxonomic and functional discrepancies between human and mouse gut microbiota, shedding light on how distinct dietary patterns may drive these differences. We observed a notable presence of taxa in the human gut microbiota that specialize in utilizing HMOs, also known as human milk glycans, reflecting specialized adaptations to the human diet.

While reinforcing findings from previous studies about the shared functional capabilities between human and mouse microbiota, our research goes further to reveal significant functional divergence between them. For instance, we discovered a pronounced enrichment of microbial protein families that metabolize food additives and preservatives within the human gut microbiota. This suggests that the human diet, rich in processed foods, may have fostered the proliferation of these metabolizing taxa. Moreover, our study illuminates the high degree of species-specificity within these microbial protein families, supporting our hypothesis that the host diet not only affects but also facilitates functional adaptation to the specific gut environment. This leads to a divergence in the taxonomic composition between humans and mice.

The MRGM has some limitations. Firstly, it includes only prokaryotic genomes, even though the importance of symbiotic viruses and fungi in animals has been increasingly recognized. Future updates should incorporate genomes for commensal viruses and fungi to create a cross-kingdom catalog of the mouse gut microbiome. Secondly, while the MRGM contains only near-complete genomes (completeness ≥ 90%), it is limited in studying certain genome regions, such as highly conserved sequences (e.g., rRNAs) and mobile sequences (e.g., genomic islands), which are notoriously difficult to assemble using short-read sequencing. Advances in long-read sequencing technology may help overcome this limitation in the near future. Thirdly, among the 1,524 species in the MRGM catalog, 404 have only a single reference genome. Given that MRGM contains only high-quality genomes (completeness ≥ 90% and contamination ≤ 5%), it is unlikely that they represent falsely assembled genomes for non-existent bacterial species. We hypothesize that the probability of assembling high-quality bacterial genomes for species with low abundance and prevalence is low. Supporting this hypothesis, we found that the number of conspecific genomes correlates with the estimated abundance and prevalence in the mouse gut microbiome (**Supplementary Fig. 1d**). This suggests that we may discover more conspecific genomes for the 404 species by analyzing additional samples with increased sequencing depth in the future. Finally, the current MRGM predominantly represents the gut bacterial genomes of laboratory mice. Since we have not extensively collected feces from wild mice, which likely have different gut microbiota due to varied diets and environments, the current depiction of the mouse gut microbiota might be quite limited. Future studies should aim to expand the catalog of mouse gut microbial genomes by including fecal samples from wild mice.

Overall, our research underscores the value of high-quality microbial genome catalogs. Such comprehensive catalogs are instrumental in enhancing our comprehension of microbial co-evolution with their hosts and the consequent shaping of gut environments and microbiome diversity, providing a clearer picture of the intricate relationship between diet, microbial function, and taxonomic diversity.

## Acknowledgements

This research was supported by the National Research Foundation of Korea (NRF) funded by the Ministry of Science and ICT (2018R1A5A2025079, 2022M3A9F3016364, 2022R1A2C1092062), and the Ministry of Trade Industry & Energy (200022947) and the Brain Korea 21 (BK21) FOUR program. P.B. was supported by the National Institute of Diabetes and Digestive and Kidney Diseases (R01 DK125382).

## Author contributions

N.K. and I.L. conceived and designed the study. P.B. and I.L. supervised the study. N.K. constructed the MRGM catalog and conducted the bioinformatic analyses. C.Y.K. provided technical guidance for MAG assembly and analysis. J.M. supplied the updated HRGM for comparative analysis. S.Y. assisted the development of the MRGM database. D.P. and S.H. generated metagenomics data. N.K. and I.L. wrote the manuscript. N.K., P.B., and I.L. edit the manuscript. The authors read and approved the final manuscript.

## References

1. Lynch SV, Pedersen O. The Human Intestinal Microbiome in Health and Disease. N Engl J Med 2016; 375:2369–79.

2. Rajpoot M, Sharma AK, Sharma A, Gupta GK. Understanding the microbiome: Emerging biomarkers for exploiting the microbiota for personalized medicine against cancer. Semin Cancer Biol 2018; 52:1–8.

3. Franzosa EA, Sirota-Madi A, Avila-Pacheco J, Fornelos N, Haiser HJ, Reinker S, et al. Gut microbiome structure and metabolic activity in inflammatory bowel disease. Nat Microbiol 2019; 4:293–305.

4. Wirbel J, Pyl PT, Kartal E, Zych K, Kashani A, Milanese A, et al. Meta-analysis of fecal metagenomes reveals global microbial signatures that are specific for colorectal cancer. Nat Med 2019; 25:679–89.

5. Nayfach S, Shi ZJ, Seshadri R, Pollard KS, Kyrpides NC. New insights from uncultivated genomes of the global human gut microbiome. Nature 2019; 568:505–10.

6. Almeida A, Mitchell AL, Boland M, Forster SC, Gloor GB, Tarkowska A, et al. A new genomic blueprint of the human gut microbiota. Nature 2019; 568:499–504.

7. Pasolli E, Asnicar F, Manara S, Zolfo M, Karcher N, Armanini F, et al. Extensive Unexplored Human Microbiome Diversity Revealed by Over 150,000 Genomes from Metagenomes Spanning Age, Geography, and Lifestyle. Cell 2019; 176:649–62 e20.

8. Almeida A, Nayfach S, Boland M, Strozzi F, Beracochea M, Shi ZJ, et al. A unified catalog of 204,938 reference genomes from the human gut microbiome. Nat Biotechnol 2021; 39:105–14.

9. Kim CY, Lee M, Yang S, Kim K, Yong D, Kim HR, et al. Human reference gut microbiome catalog including newly assembled genomes from under-represented Asian metagenomes. Genome Med 2021; 13:134.

10. Hiseni P, Rudi K, Wilson RC, Hegge FT, Snipen L. HumGut: a comprehensive human gut prokaryotic genomes collection filtered by metagenome data. Microbiome 2021; 9:165.

11. Nguyen TL, Vieira-Silva S, Liston A, Raes J. How informative is the mouse for human gut microbiota research? Dis Model Mech 2015; 8:1–16.

12. Hugenholtz F, de Vos WM. Mouse models for human intestinal microbiota research: a critical evaluation. Cell Mol Life Sci 2018; 75:149–60.

13. Xiao L, Feng Q, Liang S, Sonne SB, Xia Z, Qiu X, et al. A catalog of the mouse gut metagenome. Nat Biotechnol 2015; 33:1103–8.

14. Lesker TR, Durairaj AC, Galvez EJC, Lagkouvardos I, Baines JF, Clavel T, et al. An Integrated Metagenome Catalog Reveals New Insights into the Murine Gut Microbiome. Cell Rep 2020; 30:2909–22 e6.

15. Beresford-Jones BS, Forster SC, Stares MD, Notley G, Viciani E, Browne HP, et al. The Mouse Gastrointestinal Bacteria Catalogue enables translation between the mouse and human gut microbiotas via functional mapping. Cell Host Microbe 2022; 30:124–38 e8.

16. Kieser S, Zdobnov EM, Trajkovski M. Comprehensive mouse microbiota genome catalog reveals major difference to its human counterpart. PLoS Comput Biol 2022; 18:e1009947.

17. Li T, Yin Y. Critical assessment of pan-genomic analysis of metagenome-assembled genomes. Brief Bioinform 2022; 23.

18. Bolger AM, Lohse M, Usadel B. Trimmomatic: a flexible trimmer for Illumina sequence data. Bioinformatics 2014; 30:2114–20.

19. Langmead B, Salzberg SL. Fast gapped-read alignment with Bowtie 2. Nat Methods 2012; 9:357–9.

20. Nurk S, Meleshko D, Korobeynikov A, Pevzner PA. metaSPAdes: a new versatile metagenomic assembler. Genome Res 2017; 27:824–34.

21. Li D, Liu CM, Luo R, Sadakane K, Lam TW. MEGAHIT: an ultra-fast single-node solution for large and complex metagenomics assembly via succinct de Bruijn graph. Bioinformatics 2015; 31:1674–6.

22. Kang DD, Li F, Kirton E, Thomas A, Egan R, An H, et al. MetaBAT 2: an adaptive binning algorithm for robust and efficient genome reconstruction from metagenome assemblies. PeerJ 2019; 7:e7359.

23. Wu YW, Simmons BA, Singer SW. MaxBin 2.0: an automated binning algorithm to recover genomes from multiple metagenomic datasets. Bioinformatics 2016; 32:605–7.

24. Alneberg J, Bjarnason BS, de Bruijn I, Schirmer M, Quick J, Ijaz UZ, et al. Binning metagenomic contigs by coverage and composition. Nat Methods 2014; 11:1144–6.

25. Uritskiy GV, DiRuggiero J, Taylor J. MetaWRAP-a flexible pipeline for genome-resolved metagenomic data analysis. Microbiome 2018; 6:158.

26. Chklovski A, Parks DH, Woodcroft BJ, Tyson GW. CheckM2: a rapid, scalable and accurate tool for assessing microbial genome quality using machine learning. Nat Methods 2023; 20:1203–12.

27. Parks DH, Imelfort M, Skennerton CT, Hugenholtz P, Tyson GW. CheckM: assessing the quality of microbial genomes recovered from isolates, single cells, and metagenomes. Genome Res 2015; 25:1043–55.

28. Orakov A, Fullam A, Coelho LP, Khedkar S, Szklarczyk D, Mende DR, et al. GUNC: detection of chimerism and contamination in prokaryotic genomes. Genome Biol 2021; 22:178.

29. Afrizal A, Jennings SAV, Hitch TCA, Riedel T, Basic M, Panyot A, et al. Enhanced cultured diversity of the mouse gut microbiota enables custom-made synthetic communities. Cell Host Microbe 2022; 30:1630–45 e25.

30. Lyalina S, Stepanauskas R, Wu F, Sanjabi S, Pollard KS. Single cell genome sequencing of laboratory mouse microbiota improves taxonomic and functional resolution of this model microbial community. PLoS One 2022; 17:e0261795.

31. Ondov BD, Treangen TJ, Melsted P, Mallonee AB, Bergman NH, Koren S, et al. Mash: fast genome and metagenome distance estimation using MinHash. Genome Biol 2016; 17:132.

32. Chaumeil PA, Mussig AJ, Hugenholtz P, Parks DH. GTDB-Tk: a toolkit to classify genomes with the Genome Taxonomy Database. Bioinformatics 2019; 36:1925–7.

33. Olm MR, Brown CT, Brooks B, Banfield JF. dRep: a tool for fast and accurate genomic comparisons that enables improved genome recovery from metagenomes through de-replication. ISME J 2017; 11:2864–8.

34. Jain C, Rodriguez RL, Phillippy AM, Konstantinidis KT, Aluru S. High throughput ANI analysis of 90K prokaryotic genomes reveals clear species boundaries. Nat Commun 2018; 9:5114.

35. Saheb Kashaf S, Almeida A, Segre JA, Finn RD. Recovering prokaryotic genomes from host-associated, short-read shotgun metagenomic sequencing data. Nat Protoc 2021; 16:2520–41.

36. Parks DH, Chuvochina M, Rinke C, Mussig AJ, Chaumeil PA, Hugenholtz P. GTDB: an ongoing census of bacterial and archaeal diversity through a phylogenetically consistent, rank normalized and complete genome-based taxonomy. Nucleic Acids Res 2022; 50:D785–D94.

37. Nguyen LT, Schmidt HA, von Haeseler A, Minh BQ. IQ-TREE: a fast and effective stochastic algorithm for estimating maximum-likelihood phylogenies. Mol Biol Evol 2015; 32:268–74.

38. Letunic I, Bork P. Interactive Tree Of Life (iTOL) v5: an online tool for phylogenetic tree display and annotation. Nucleic Acids Res 2021; 49:W293–W6.

39. Seemann T. barrnap 0.9 : rapid ribosomal RNA prediction https://github.com/tseemann/barrnap. Github, 2018.

40. Wood DE, Lu J, Langmead B. Improved metagenomic analysis with Kraken 2. Genome Biol 2019; 20:257.

41. Lu J BF, Thielen P, Salzberg SL. Bracken: estimating species abundance in metagenomics data. PeerJ Computer Science 2017.

42. Seemann T. Prokka: rapid prokaryotic genome annotation. Bioinformatics 2014; 30:2068–9.

43. Steinegger M, Soding J. Clustering huge protein sequence sets in linear time. Nat Commun 2018; 9:2542.

44. Cantalapiedra CP, Hernandez-Plaza A, Letunic I, Bork P, Huerta-Cepas J. eggNOG-mapper v2: Functional Annotation, Orthology Assignments, and Domain Prediction at the Metagenomic Scale. Mol Biol Evol 2021; 38:5825–9.

45. Huerta-Cepas J, Szklarczyk D, Heller D, Hernandez-Plaza A, Forslund SK, Cook H, et al. eggNOG 5.0: a hierarchical, functionally and phylogenetically annotated orthology resource based on 5090 organisms and 2502 viruses. Nucleic Acids Res 2019; 47:D309–D14.

46. Mistry J, Chuguransky S, Williams L, Qureshi M, Salazar GA, Sonnhammer ELL, et al. Pfam: The protein families database in 2021. Nucleic Acids Res 2021; 49:D412–D9.

47. Kanehisa M, Furumichi M, Sato Y, Kawashima M, Ishiguro-Watanabe M. KEGG for taxonomy-based analysis of pathways and genomes. Nucleic Acids Res 2023; 51:D587–D92.

48. Gene Ontology C. The Gene Ontology resource: enriching a GOld mine. Nucleic Acids Res 2021; 49:D325–D34.

49. Drula E, Garron ML, Dogan S, Lombard V, Henrissat B, Terrapon N. The carbohydrate-active enzyme database: functions and literature. Nucleic Acids Res 2022; 50:D571–D7.

50. Bairoch A. The ENZYME database in 2000. Nucleic Acids Res 2000; 28:304–5.

51. Kulmanov M, Hoehndorf R. DeepGOPlus: improved protein function prediction from sequence. Bioinformatics 2020; 36:422–9.

52. Suzek BE, Wang Y, Huang H, McGarvey PB, Wu CH, UniProt C. UniRef clusters: a comprehensive and scalable alternative for improving sequence similarity searches. Bioinformatics 2015; 31:926–32.

53. Tonkin-Hill G, MacAlasdair N, Ruis C, Weimann A, Horesh G, Lees JA, et al. Producing polished prokaryotic pangenomes with the Panaroo pipeline. Genome Biol 2020; 21:180.

54. Hsieh TC, Ma KH, Chao A. iNEXT: an R package for rarefaction and extrapolation of species diversity (Hill numbers). Methods Ecol Evol 2016; 7:1451–6.

55. Manghi P, Blanco-Miguez A, Manara S, NabiNejad A, Cumbo F, Beghini F, et al. MetaPhlAn 4 profiling of unknown species-level genome bins improves the characterization of diet-associated microbiome changes in mice. Cell Rep 2023; 42:112464.

56. Fritz A, Hofmann P, Majda S, Dahms E, Droge J, Fiedler J, et al. CAMISIM: simulating metagenomes and microbial communities. Microbiome 2019; 7:17.

57. Campbell C, Dikiy S, Bhattarai SK, Chinen T, Matheis F, Calafiore M, et al. Extrathymically Generated Regulatory T Cells Establish a Niche for Intestinal Border-Dwelling Bacteria and Affect Physiologic Metabolite Balance. Immunity 2018; 48:1245–57 e9.

58. Pasolli E, Truong DT, Malik F, Waldron L, Segata N. Machine Learning Meta-analysis of Large Metagenomic Datasets: Tools and Biological Insights. PLoS Comput Biol 2016; 12:e1004977.

59. Thomas AM, Manghi P, Asnicar F, Pasolli E, Armanini F, Zolfo M, et al. Metagenomic analysis of colorectal cancer datasets identifies cross-cohort microbial diagnostic signatures and a link with choline degradation. Nat Med 2019; 25:667–78.

60. Blin K, Shaw S, Kloosterman AM, Charlop-Powers Z, van Wezel GP, Medema MH, et al. antiSMASH 6.0: improving cluster detection and comparison capabilities. Nucleic Acids Res 2021; 49:W29–W35.

61. Kautsar SA, van der Hooft JJJ, de Ridder D, Medema MH. BiG-SLiCE: A highly scalable tool maps the diversity of 1.2 million biosynthetic gene clusters. Gigascience 2021; 10.

62. Kim CY, Ma J, Lee I. HiFi metagenomic sequencing enables assembly of accurate and complete genomes from human gut microbiota. Nat Commun 2022; 13:6367.

63. Miller CS, Baker BJ, Thomas BC, Singer SW, Banfield JF. EMIRGE: reconstruction of full-length ribosomal genes from microbial community short read sequencing data. Genome Biol 2011; 12:R44.

64. Galperin MY, Wolf YI, Makarova KS, Vera Alvarez R, Landsman D, Koonin EV. COG database update: focus on microbial diversity, model organisms, and widespread pathogens. Nucleic Acids Res 2021; 49:D274–D81.

65. Eisenhofer R, Odriozola I, Alberdi A. Impact of microbial genome completeness on metagenomic functional inference. ISME Commun 2023; 3:12.

66. Malmstrom RR. Quality MAGnified. Nat Rev Microbiol 2023.

67. Sharon I, Morowitz MJ, Thomas BC, Costello EK, Relman DA, Banfield JF. Time series community genomics analysis reveals rapid shifts in bacterial species, strains, and phage during infant gut colonization. Genome Res 2013; 23:111–20.

68. Scholz M, Ward DV, Pasolli E, Tolio T, Zolfo M, Asnicar F, et al. Strain-level microbial epidemiology and population genomics from shotgun metagenomics. Nat Methods 2016; 13:435–8.

69. De Filippis F, Pasolli E, Tett A, Tarallo S, Naccarati A, De Angelis M, et al. Distinct Genetic and Functional Traits of Human Intestinal Prevotella copri Strains Are Associated with Different Habitual Diets. Cell Host Microbe 2019; 25:444–53 e3.

70. Zhao S, Lieberman TD, Poyet M, Kauffman KM, Gibbons SM, Groussin M, et al. Adaptive Evolution within Gut Microbiomes of Healthy People. Cell Host Microbe 2019; 25:656–67 e8.

71. Tamames J, Cobo-Simon M, Puente-Sanchez F. Assessing the performance of different approaches for functional and taxonomic annotation of metagenomes. BMC Genomics 2019; 20:960.

72. Blanco-Miguez A, Beghini F, Cumbo F, McIver LJ, Thompson KN, Zolfo M, et al. Extending and improving metagenomic taxonomic profiling with uncharacterized species using MetaPhlAn 4. Nat Biotechnol 2023.

73. Ye SH, Siddle KJ, Park DJ, Sabeti PC. Benchmarking Metagenomics Tools for Taxonomic Classification. Cell 2019; 178:779–94.

74. Sun Z, Huang S, Zhang M, Zhu Q, Haiminen N, Carrieri AP, et al. Challenges in benchmarking metagenomic profilers. Nat Methods 2021; 18:618–26.

75. Wright RJ, Comeau AM, Langille MGI. From defaults to databases: parameter and database choice dramatically impact the performance of metagenomic taxonomic classification tools. Microb Genom 2023; 9.

76. Beghini F, McIver LJ, Blanco-Miguez A, Dubois L, Asnicar F, Maharjan S, et al. Integrating taxonomic, functional, and strain-level profiling of diverse microbial communities with bioBakery 3. Elife 2021; 10.

77. Cani PD, Depommier C, Derrien M, Everard A, de Vos WM. Akkermansia muciniphila: paradigm for next-generation beneficial microorganisms. Nat Rev Gastroenterol Hepatol 2022; 19:625–37.

78. Brennan CA, Garrett WS. Fusobacterium nucleatum - symbiont, opportunist and oncobacterium. Nat Rev Microbiol 2019; 17:156–66.

79. Vacca M, Celano G, Calabrese FM, Portincasa P, Gobbetti M, De Angelis M. The Controversial Role of Human Gut Lachnospiraceae. Microorganisms 2020; 8.

80. Gavzy SJ, Kensiski A, Lee ZL, Mongodin EF, Ma B, Bromberg JS. Bifidobacterium mechanisms of immune modulation and tolerance. Gut Microbes 2023; 15:2291164.

81. Turroni F, Milani C, Duranti S, Mahony J, van Sinderen D, Ventura M. Glycan Utilization and Cross-Feeding Activities by Bifidobacteria. Trends Microbiol 2018; 26:339–50.

82. De Angelis M, Ferrocino I, Calabrese FM, De Filippis F, Cavallo N, Siragusa S, et al. Diet influences the functions of the human intestinal microbiome. Sci Rep 2020; 10:4247.

83. Peng QN, Huo DX, Ma CC, Jiang SM, Wang LS, Zhang JC. Monosodium glutamate induces limited modulation in gut microbiota. J Funct Foods 2018; 49:493–500.

84. Heilbronner S, Krismer B, Brotz-Oesterhelt H, Peschel A. The microbiome-shaping roles of bacteriocins. Nat Rev Microbiol 2021; 19:726–39.

85. Arun KB, Sindhu R, Alex D, Binod P, Pughazhendi A, Joseph TC, et al. Bacterial bioactive metabolites as therapeutic agents: From production to action. Sustain Chem Pharm 2022; 27.

