## Supplementary Figure 1 for "MRGM: An enhanced catalog of mouse gut microbial genomes substantially broadening taxonomic and functional landscapes"

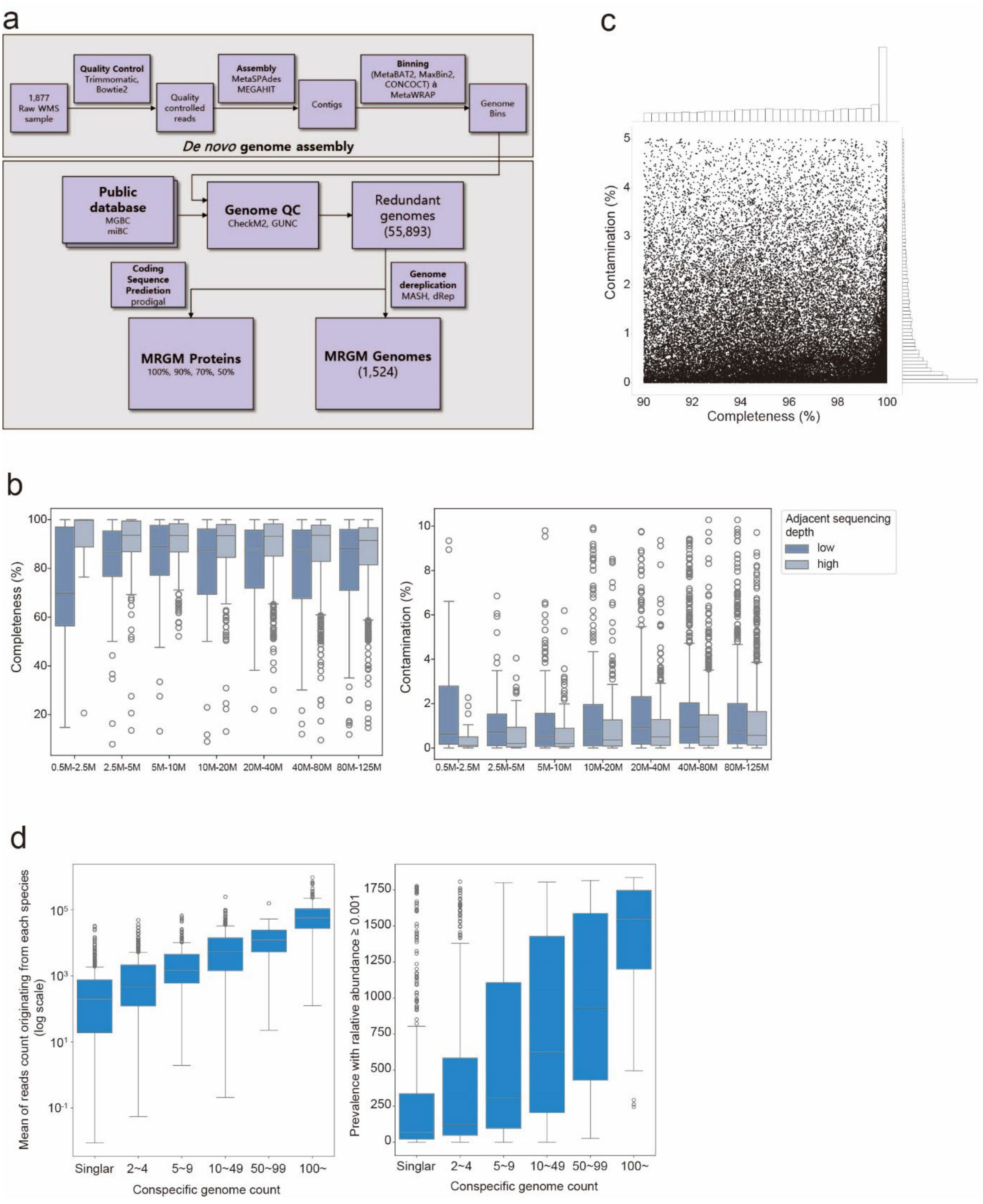


**Supplementary Figure 1. Summary of MRGM construction.** (**a**) Summary of in-house genome-resolved metagenomic analysis pipeline used for the MRGM construction. (**b**) Box plots showing the completeness (left) and contamination (right) of simulated datasets using 10 metagenome samples analyzed by ultra-deep sequencing. For random samples of 0.5, 2.5, 5, 10, 20, 40, 80, and 125 million read pairs, we assembled MAGs for 80 simulated datasets (10 samples × 8 depths). The quality of MAGs (completeness and contamination) was compared for the same species (with Mash ANI ≥ 90%) across adjacent sequencing depths. (**c**) Summary of the completeness and contamination of 42,245 non-redundant genomes of the MRGM. (**d**) Box plots showing the read count for each species as an estimate of species abundance (left) and the number of detected samples for each species, determined by a relative abundance ≥ 0.001, as an estimate of species prevalence (right) for each group of species within the specified range of conspecific genomes.
